# Investigating the Consequences of Non-active site Mutations on the Structure, Function and Dynamics of the Molten Globule Enzyme Monomeric Chorismate Mutase

**DOI:** 10.64898/2026.04.11.717874

**Authors:** Shrutidhara Biswas, Pamarthi Gangadhar, Srihari Pabbaraja, Rajaram Swaminathan

**Affiliations:** Department of Biosciences and Bioengineering, Indian Institute of Technology Guwahati, Guwahati 781039, Assam, India; Department of Organic Synthesis and Process Chemistry, CSIR-Indian Institute of Chemical Technology, Tarnaka, Hyderabad 500007, Telangana, India; Academy of Scientific and Innovative Research (AcSIR), Ghaziabad 201002, India

## Abstract

Intrinsically disordered enzymes serve as useful models to understand their catalytic function against the backdrop of an unstructured protein. The characteristic flexibility in conformation seen in IDPs is a rare occurrence among enzymes and one such enzyme is the engineered protein: monomeric Chorismate Mutase (mCM). mCM surprisingly retains similar enzyme activity as its parent dimeric protein Chorismate Mutase from *Methanococcus jannaschii* (MjCM) despite losing the ordered globular structure. In this work using a previously demonstrated transition state analogue (TSA), we analyze the structural transitions in mCM during catalysis. Additionally, consequences of three non-active site single point mutations were investigated using CD; Trp-Dansyl FRET measurements using fluorescence lifetime; and time-resolved fluorescence anisotropy measurements; to map the local (near Trp) and global structural transitions in mCM during catalysis. Mutant2 (W24K + C69A); and Mutant3 (W24K + C69A + A6C); revealed a 97 and 89% drop-in activity compared to mCM; quite unlike Mutant1 (W24K, 19% drop). Mutant1 as opposed to Mutant3 was most sensitive to binding of TSA as quantified by structural displacement measured using FRET. This was consistent with an overall globular structure compaction induced by TSA binding in Mutant1 as reflected by a dip in rotational correlation time of Cys-conjugated dansyl probe from 10.3 to 8.4 ns. Our results highlight the critical role of Cys69 residue, that is ~19 Å away from mCM active site, in influencing the hydrophobic collapse upon substrate binding and subsequent catalytic activity.

## INTRODUCTION

Intrinsically disordered proteins (IDPs) are a class of proteins gaining immense attention over the last two decades due to their unique structural characteristics and relevance in many major human diseases [1,2,3]. They are known to carry out a variety of crucial biological functions viz. signaling, regulation and transcription despite lacking a well-defined tertiary structure [4,5]. The discovery that a native IDP exists as ensembles of inter-converting highly flexible protein structures that can adopt an ordered conformation when bound to appropriate ligand(s) revolutionized the way protein structure-function interactions are interpreted [6]. An interesting subclass within this IDP family is the ID enzyme. Such enzymes can carry out their function without a rigid catalytic core, previously thought to be a pre-requisite for efficient enzyme catalysis [7,9]. UreG, a GTPase, was the first discovered ID enzyme [9] and till date a large repertoire of enzymes with ID regions have been identified [10].

The role of ID regions in many enzymes has been found to be associated in modulation of its specificity or in self-regulation [10]. But cases of ID enzymes where the entire protein assumes a disordered state, such as a molten globule, is very rare and only few such cases have been reported [11,12,13,14]. Molten globules are closely packed ensemble of conformations with intact secondary structure but negligible tertiary structure. They are mostly studied as intermediates in protein folding which may possess some reminiscent function of a native protein [15,16].

Monomeric chorismate mutase (mCM), is one such disordered molten globule enzyme but with ability to function almost identical to the native ordered dimeric CM enzyme [17,18]. mCM can convert its substrate chorismate to prephenate with a *k*_*cat*_ of 3.2 s^−1^ and *K*_*m*_ of 170 μM; values comparable to the parent CM enzyme. Surprisingly, mCM also retains a high degree of global structural flexibility even after substrate binding [18]. The unusual coupling of such flexibility and functionality in an enzyme inspired us to carry out a meticulous scrutiny of the mCM structural transitions and function associated dynamics.

The simple four helical bundle structure, small size (12.74 kDa, 104 amino acid residues) and the presence of only two tryptophans and a single cysteine residue allowed us to investigate mCM through fluorescence based experimental methods without many limitations. We probed the structure and dynamics of mCM using the intrinsic fluorophore tryptophan and the extrinsic fluorophore dansyl (labeled to the sole cysteine residue), which also served as a FRET pair in our experiments. Through site-directed mutagenesis, single tryptophan and single cysteine mutant pair were generated at specific locations in mCM protein using available NMR structure (PDB: 2GTV) with substrate analogue (TSA). This enabled distance between a Trp (donor)— Dansyl(acceptor) pair to be measured effectively using FRET, without perturbing the substrate binding site in the enzyme. FRET which acts as a molecular ruler [19,20] was used to measure structural transitions in mCM in terms of change in proximity between indole (trp) and dansyl (cys) groups, *associated with binding of transition state analogue* (TSA) in mCM. In addition to FRET, nanosecond rotational motion of Cys-conjugated dansyl probe was measured using time-resolved fluorescence anisotropy to obtain quantitative information on the Brownian rotational diffusion of dansyl group at the local and global (whole protein) scales in presence and absence of TSA. Finally, the enzyme kinetic parameters of mCM and mutant proteins were determined, so as to connect the structural information gleaned from FRET and fluorescence anisotropy with functional observations.

For this work, three mutants namely Mutant1 (W24K); Mutant2 (W24K/C69A); and Mutant3 (W24K/C69A/A6C); were generated to enable investigations as detailed above. The structural and functional characteristics of the mutants obtained from fluorescence and enzyme experiments were compared with those for parent mCM enzyme with the objective of obtaining molecular insights on how the mCM secondary/tertiary structure and folding/packing are coupled with enzymatic activity.

## EXPERIMENTAL METHODS

### Generation of Mutants

Selection for mutation sites and the generation of mutant proteins were based on several considerations: (1) Appropriate distance for efficient FRET; (2) Picking residues that are not part of the active site of the enzyme; (3) Mutant 1 (W24K) was chosen for two reasons: a) to simplify data analysis from the single Trp26 and single Cys69 or Cys6 (Mutant 3) in the FRET pair, b) Mutant 1 has been previously shown to be active [17]; (4) Replacement of Trp and Cys was done with residues that inflict minimal structural perturbations (Lys and Ala respectively); (5) Mutant 3 served as an alternative Trp/Cys pair to compare with Mutant 1, with Cys positioned farther away; (6) Mutant2, an intermediate product during mutation (no Cys), was studied to understand the role of Cys69 in mCM. In accordance with our mutation plans, primers were designed using the Agilent QuikChange Primer designing software. All the primers were purchased from Integrated DNA Technologies, Inc. The parent DNA plasmid (mMjCM) used as the template for mutagenesis was generously gifted by Prof. Donald Hilvert’s lab in ETH Zurich, Switzerland. The mutations were carried out using the QuikChange II Site-directed Mutagenesis Kit (200524). Correct incorporation of mutations in all variants of the parent plasmid were confirmed by DNA sequencing.

### Purification of mCM and mutants

The plasmids were transformed into *E. coli* BL21 (DE3) cells and used for over expression of all mCM variants. Bacterial cultures were grown in LB media, containing 100 µg/mL ampicillin (at 37 °C, 180 rpm) and induced with Isopropyl β-D-1-thiogalactopyranoside (IPTG, 1 mM). Cells were pelleted and then resuspended in ice cold lysis buffer (20 mM Tris.HCl pH 7.4; 150 mM NaCl; 0.5% Triton-X; 1% glycerol; 1 mM PMSF). Resuspended cells were disrupted by sonication and centrifuged to separate debris. Supernatants were filtered with a 0.45 µm filter (Millipore) and then allowed to bind to a pre-equilibrated (20 mM Tris.HCl pH 7.4; 150 mM NaCl; 20 mM Imidazole) Ni Sephrose HisTrap column (GE). The supernatant was circulated through the column using a peristaltic pump (GE Healthcare) to allow protein binding. Unbound protein was washed out with a wash buffer (20 mM Tris.HCl pH 7.4; 150 mM NaCl; 75 mM Imidazole). The bound protein was eluted with an elution buffer (20 mM Tris.HCl pH 7.4; 150 mM NaCl; 350 mM Imidazole). Purity of purified proteins checked on SDS PAGE and further analyzed by mass spectrometry using a MALDI-TOF mass spectrometer (Bruker Daltonics, Germany).

### Enzyme Assay

To measure the enzymatic activity of mCM, the disappearance of its substrate (Chorismic acid) was monitored spectroscopically at 274 nm (ε_274_ = 2630 M^−1^cm^−1^). Readings were recorded in a UV visible spectrophotometer (Perkin Elmer) at every 1 s interval for 300 s. 150 nM of mCM enzyme in PBS buffer pH 7.2 supplemented with BSA (0.1 mg/ml) was used for the assay. A varying substrate concentration of 50-400 µM was utilized for the enzymatic reaction. The data obtained is extrapolated in the Eadie-Hofstee plot to obtain the *K*_*m*_, *V*_*max*_ and *k*_*cat*_ values.

### mCM labelling with dansyl probe

Labeling of single cysteine in mCM, Mutant 1 and Mutant 3 with the IAEDANS (1,5-IAEDANS, 5-((((2-iodoacetyl)amino)ethyl)amino)naphthalene-1-sulphonic acid) was carried out as recommended by Molecular Probes with slight modifications [21]. 100 μL of 100 mM IAEDANS was added slowly to 900 μL of 100 µM mCM (20 mM Tris.HCl, pH 9;150 mM NaCl) to make a total reaction volume of 1 ml. This reaction mixture was incubated at room temperature (25 °C) for three hours with continuous stirring at 150 rpm. Free dye was removed from the dansyl labeled protein by passing through pre-equilibrated PD-10 desalting column (20 mM Tris.HCl pH 9; 150 mM NaCl). The labeled protein was eluted from the column in PBS buffer pH 7.4. Concentration of protein and dye was measured by using absorbance at 280 and 337 nm, respectively. The molar extinction coefficient of conjugated probe with protein at 337 nm is 6100 M^−1^cm^−1^ [22]. The dye to protein labelling ratio was determined to be 0.9:1.

### TSA Synthesis

The Transition State Analogue (TSA), (8-hydroxy-2-oxa-bicyclo[3.3.1]non-6-ene-3,5-dicarboxylic acid), was synthesized by Dr. P. Gangadhar in Prof. P. Srihari’s lab (IICT Hyderabad, India). The synthesis process was accomplished by using Bartlett’s published protocol [23]. More details on the methodology, NMR characterization of intermediate and final products synthesized are shown in Supplementary Information (see last section).

### UV-Visible absorption spectra

The absorption spectrum of all mCM variants was recorded using double beam Lambda-25 UV-Vis Spectrophotometer (Perkin Elmer, USA) at room temperature (25 °C). Extinction coefficients (ε) at λ_280_ nm 15,595 M^−1^cm^−1^ for mCM; 10,095 M^−1^cm^−1^ for Mutant1 and Mutant3; 9975 M^−1^cm^−1^ for Mutant2; were used for the estimation of protein concentration using the Beer-Lambert equation.

### Circular Dichroism measurements

CD spectra of mCM and mutants were recorded in a CD spectrometer (Make: Jasco, Model J-1500, Jasco Inc., Maryland, USA), equipped with a Peltier thermoregulator. Protein samples were prepared by dissolving 10 μM of mCM in 10 mM phosphate buffer supplemented with 10 mM NaF pH 7.4. All CD spectra were collected from 190 to 260 nm, at a scan speed of 100 nm/min, bandwidth of 2 nm and data pitch of 0.1 nm. All spectra were collected using 2 mm quartz cuvette (Starna Scientific Ltd). CD spectra were analyzed using K2D3 server and a useful estimate of the α-helical content and its variation across the mCM proteins was made using this analysis.

Chemical denaturation of mCM and its mutants were performed by incubating protein samples in increasing concentration of GdnHCl (0-6 M). Changes in the secondary structure were monitored by following the changes in ellipticity at 222 nm at 25 °C. Data points obtained were fitted by non-linear regression analysis (dose-response curve) using OriginPro 8 software. For thermal denaturation, change in ellipticity of mCM and the mutants were continuously monitored at 222 nm in the temperature range from 25 to 90 °C and reverse with a heating/cooling rate of 1 °C/min. CD spectral scans from 260 nm to 190 nm at specific temperature intervals and their reverse scans were also recorded during the experiment.

### Steady state fluorescence and anisotropy measurements

All measurements were performed using Fluoromax-4 Spectrofluorometer (Jobin-Yvon Horiba Inc., USA) equipped with motorized polarizers. For all mCM Trp steady state fluorescence intensity measurements, samples were excited at 295 nm and emission was collected between 310 nm to 500 nm. For acquiring steady state fluorescence intensity of dansyl labeled at cysteine of mCM, all samples were excited at 340 nm and emission collected between 355 nm to 650 nm. All steady state fluorescence measurements were repeated at least three times at room temperature (25 °C) in 200 μL quartz cuvette with a 10 mm pathlength. For Trp steady state anisotropy, all mCM samples were excited at 295 nm and emission collected at 345 nm. For steady state anisotropy of dansyl labeled with mCM, all samples were excited at 340 nm and emission was collected at 505 nm. All steady state anisotropy data were G-factor corrected and each anisotropy measurement was an average of 10 independent measurements.

### ANS binding assay

Stock of ANS (1mg/mL) was prepared in deionized water and 20 μM of mCM (dissolved in PBS pH 7.4) was mixed with 40 μM of ANS. To study the effect of TSA on ability of ANS to bind the ligand-bound mCM (all variants), 40 μM of TSA was added to 20 μM of mCM and incubated for an hour before addition of 40 μM of ANS to the sample mixture. Binding of ANS with mCM was monitored by measuring the steady state fluorescence intensity of ANS. All samples were excited at 380 nm and emission was collected between 400 nm to 700 nm. All ANS fluorescence measurements were done in triplicate at room temperature (25°C).

### Time-resolved Fluorescence and Anisotropy measurements

All fluorescence intensity decay measurements were done by time-correlated single photon counting (TCSPC) instrument, using the ‘DeltaPro’ equipped with motorized polarizer, supplied by Horiba Scientific, UK. For Trp, excitation was done with pulsed light source, 295 nm DeltaDiode™ having about 0.81 ns Instrument response function (IRF) FWHM and a 20 MHz repetition rate. Emission of Trp was collected with 320 nm longpass filter (WG320) to block the excitation light. Total peak counts were collected up to 15,000 counts and fluorescence intensity decay was collected in 2202 channels (in 0 to 61.66 ns time range) with temporal resolution of 0.028 ns/ channel. For dansyl labeled with Cys of mCM (Cys69/Cys6), the excitation was done with a pulsed 340 nm DeltaDiode light source. Emission of dansyl was measured with 370 nm longpass filter (KV370) and peak counts were measured up to 20,000 and fluorescence intensity decay was collected in 3995 channels (in 0 to 223.72 ns time range) with temporal resolution of 0.056 ns/channel. IRF was collected using scattering solution of colloidal silica, without emission filter for each of the sample.

Decay of time-dependent fluorescence intensity was expressed as a multi-exponential:

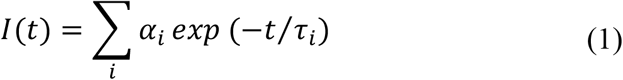

Here *τ*_*i*_ and *α*_*i*_ represent the *i*^*th*^ lifetime and *i*^*th*^ fractional amplitude, respectively for the *i*^*th*^ decay component. Sum of *α*_*i*_ is equal to unity and *i* can take values from 1—3. The mean lifetime τm can be calculated as follows:

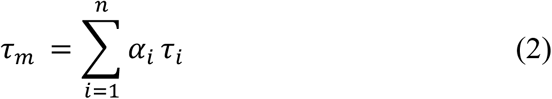

Fluorescence lifetime of Trp and dansyl conjugated with cysteine of mCM was obtained by fitting *I(t)* to Eq. 1 using non-linear least squares and iterated reconvolution employing DAS6 v6.8 (Horiba); exp2 and exp3 decay analysis software [25].

For time-resolved anisotropy measurements, the intensity decays *I*_∥_(*t*) and *I*_⊥_(*t*) representing the sample intensity decay measured with the emission polarizer in the parallel and perpendicular orientations, respectively was collected.

The time-resolved anisotropy decay data was analyzed on the basis of equations given below

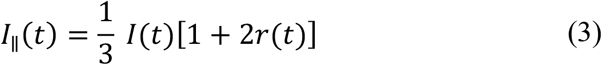

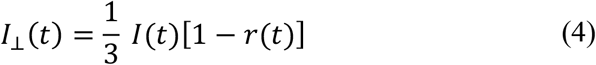

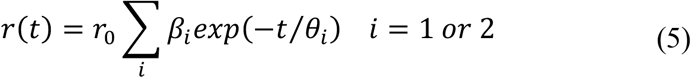

Here, *r(t)* represents the anisotropy decay, *r*_*0*_ denotes the initial anisotropy, amplitude of i^th^ rotational correlation time *θ*_*i*_ is denoted as *β*_*i*_ and *∑*_*i*_ *β*_*i*_ = 1. Rotational correlation times of dansyl probe were extracted from tani4 software by fitting *i*_∥_(*t*) and *i*_⊥_(*t*) in equations 3 and 4, globally employing non-linear least squares analysis (Marquardt method)[25]. The goodness of fit was evaluated by analyzing the reduced χ^2^ and randomness of residuals.

The values collected in the above fitting analysis were used to yield the steady-state anisotropy *r*_*ss*_ [25].

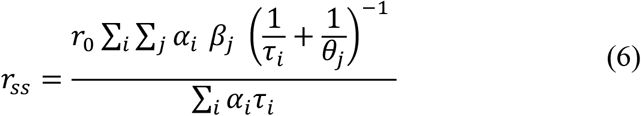

Further, the calculated steady-state anisotropy values were compared with the steady-state anisotropy values measured independently by using spectrofluorometer, to cross verify the fitted values of *θ*_*i*_, *β*_*i*_ and *r*_*0*_.

Fluorescence lifetime distributions were obtained by the maximum entropy method (MEM) as described previously [24,26]. Software for exp2; exp3; tani4 and MEM were kindly provided by Prof.. N. Periasamy, Tata Institute of Fundamental Research (TIFR), Mumbai, India.

### Intramolecular Förster Resonance Energy Transfer (FRET) measurements

To calculate the distance between indole (Trp) and dansyl group (conjugated at Cys) in different variants of mCM under various experimental conditions, fluorescence lifetime of Trp in absence of the dansyl acceptor (τ_D_), and in presence of the acceptor (τ_DA_) was monitored by TCSPC with sub-nanosecond resolution. The Trp was excited with pulsed light source, 295 nm DeltaDiode and subsequently emission of Trp was collected with 330-355 nm bandpass filter (ET340/40X) supplied by CHROMA, to avoid collecting emission light coming from the dansyl probe. Peak counts were measured up to 15,000 and fluorescence intensity decay was collected in 2202 channels (in 0 to 61.66 ns time range) with temporal resolution of 0.028 ns/channel. All lifetime measurements were done at room temperature and for each sample at least three independent lifetime measurements were recorded. Förster distance (R_0_) 22 Å, for tryptophan-IAEDANS pair was used [27] for FRET efficiency (E) calculation.

The FRET efficiency was calculated from the mean lifetime of donor (Tryptophan), in absence and presence of acceptor (Dansyl) as described below.

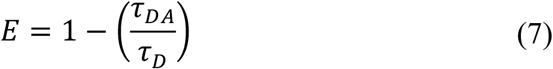

For FRET analysis, the κ value used was 0.816 (κ^2^ = 2/3) and value for η used was 1.00 cP [28].

## RESULTS

### Steady-state Trp fluorescence in mCM reveals the indole microenvironment while ANS fluorescence illuminates the global molten globule state

The fluorescence emission spectrum of mCM collected after excitation at 295 nm displayed an emission maximum at 340 nm (Figure 1A), suggesting that the Trp residues (W24, W26) are significantly exposed to the aqueous solvent. The engineered mCM has two Trp residues in close vicinity positioned on a flexible turn connecting two of its α-helices [17,29]. A similar emission profile was observed for the mutants, where the fluorescence intensities were slightly elevated and the average emission maxima red-shifted by 1-2 nm (Figure 1A). The higher fluorescence intensity in mutants was surprising as only one Trp residue was present. A probable explanation could be changes in the Trp (W26) microenvironment following the W24K mutation. Also, we have evidence of homoFRET between the two Trp residues in mCM (Figure S1) which could have quenched and affected the Trp fluorescence intensity of mCM when compared to its mutant forms. Fluorescence spectra from an excess of TSA (1mM) alone did not result in a large enough fluorescence signal that could interfere with Trp fluorescence from mCM (Figure 1, inset). Thus, it was possible to derive clear Trp fluorescence data with very low background signal in the presence of TSA. mCM Trp fluorescence in the absence of Trp-Dansyl FRET remained unaffected by any bound TSA (Figure S2), implying that (a) TSA binding does not directly impact Trp fluorescence and (b) any transition in mCM structure after ligand binding does not involve the short flexible loop containing the Trp(s). As steady-state fluorescence can be sensitive to sample concentrations, these findings were later confirmed by Trp fluorescence lifetime (Table S4).

**Figure 1.**
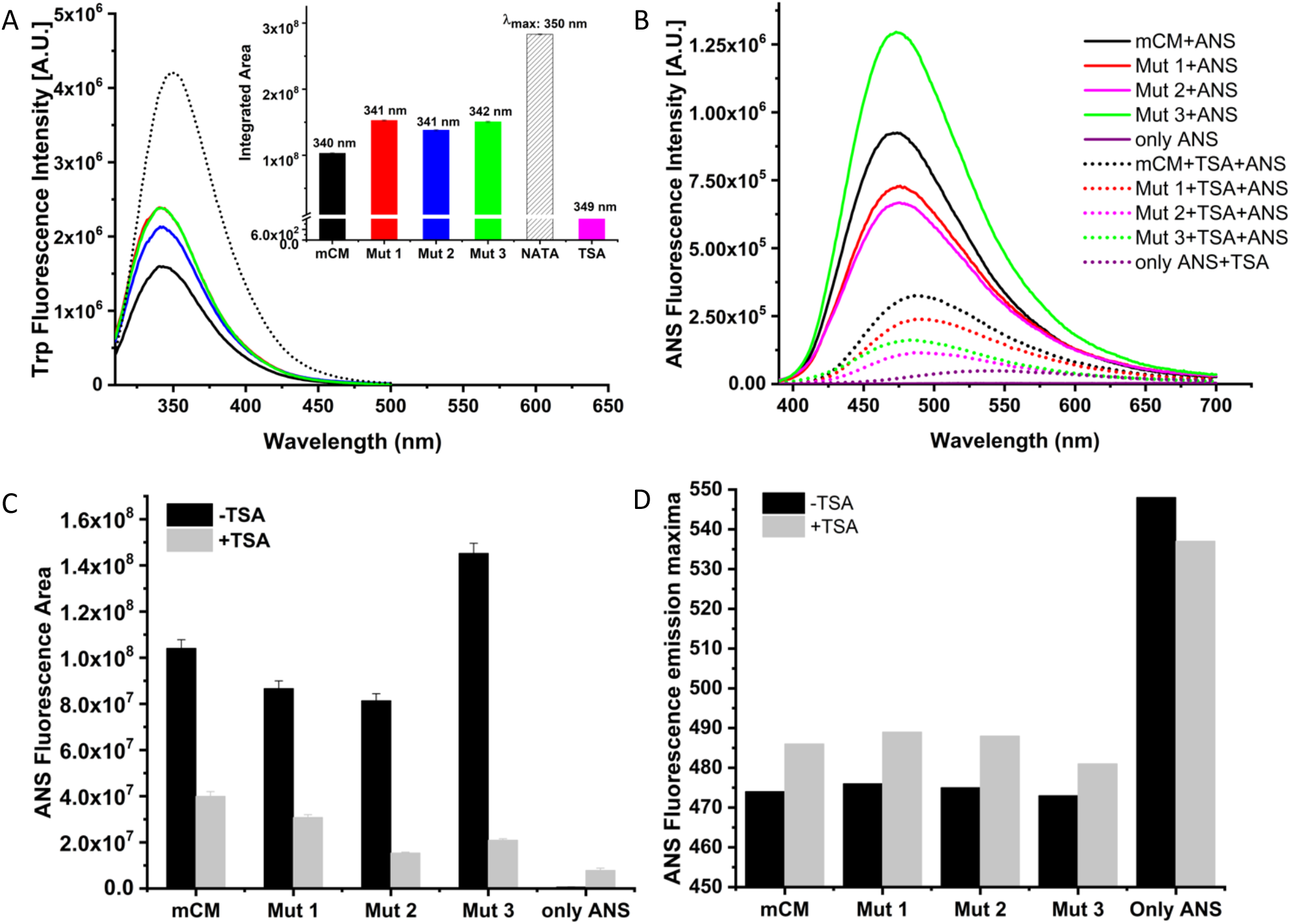
Fluorescence emission spectra of mCM and its mutants. (A) Emission spectra obtained after excitation of Trp at 295 nm. 20 μM of each of the proteins were used for the experiment. Fluorescence emission of NATA (10 μM) is shown as a dotted curve. The integrated fluorescence areas of all the curves are shown as an inset. TSA integrated area is 10^5^ times less than mCM. (B) The spectra revealing the binding of ANS to mCM and all the mutants (solid lines). Adding TSA to mCM before ANS binding (dotted lines) negatively affected the binding process. All the samples were excited at 375 nm. (C) Integrated fluorescence area and (D) emission λmax wavelengths corresponding to the ANS fluorescence intensities for mCM and its mutants in absence and presence of TSA.

ANS is known to bind preferably to molten globular states of proteins [15]. ANS binding to mCM and its mutants led to a dramatic increase in ANS fluorescence. ANS displayed a lower fluorescence intensity in Mutant1 and Mutant2 compared to mCM (Figure 1B, 1C). The hydrophobic core/patches of Mutant1 and Mutant 2 could have become more solvent exposed due to the specific point mutations, thus affecting local and/or global conformation of the protein. Surprisingly, Mutant 3 reported higher ANS fluorescence than mCM. Structural re-arrangement of the helices in this triple mutant is possible, giving rise to additional or altogether new hydrophobic pockets where ANS can bind. Furthermore, TSA was incubated with mCM and the mutants before performing the ANS assay. Interestingly, a significant reduction in intensity and red shift of emission maxima (Figure 1B, 1C, 1D) in ANS fluorescence was observed in all TSA bound proteins, though ANS could still bind mCM to some extent. This reduction in ANS binding was even more prominent in the mutants (Figure 1B, 1C). This can happen if TSA binding prevents ANS from accessing its binding sites due to structural compaction of mCM after ligand binding, which can also be aggravated by a possible competition with a more specific ligand like TSA for the hydrophobic core region. In a previous study by Vamvaca and coworkers (17), the ability of ANS to bind mCM in the TSA bound form was interpreted as a retention of the dynamic nature of mCM even after the transition to a more ordered state. In our ANS assay, the slight decrease in ANS fluorescence in all mutants compared to mCM suggests that either (a) the mutants bind TSA similarly as mCM but may have incurred some loss in their dynamic nature in the ligand bound state or (b) the mutants may have acquired relatively more compact structure after ligand binding, further limiting ANS access to the hydrophobic core. The latter conclusion is less likely as our time-resolved fluorescence anisotropy data (Table S1, S2, S3) indicates an overall decrease in global compaction in the mutants as discussed in the following section.

### Global structural compaction in mCM after ligand binding was observed using time-resolved fluorescence anisotropy decay of covalently conjugated dansyl probe

The fluorescence anisotropy decay of the dansyl probe conjugated mCM (at Cys69.SH or Cys6.SH) bound under various conditions (with/without TSA) were analysed and then compared to extract information on the local dynamics surrounding the probe as well as global tumbling rotational diffusion rate of the probe. The intrinsic Trp in mCM, having a short fluorescence lifetime (~ 2 ns), was not used for anisotropy decay studies. The ligand-free mCM (30 μM) displayed a fast rotational component of 1.32 ns (amp=0.44) and a slow rotational component of 8.6 ns (amp=0.56). The fast component corresponds to local rotational motion around the dansyl probe, whereas the slow component relates to the global rotation of the entire protein. A sharp decrease in the value of the fast component to 0.71 ns (for 1:2 ratio of mCM to TSA) and further down to 0.44 ns (for 1:4 ratio) with increased amplitude (0.44 to 0.61) suggested that the local rotational motion of the dansyl probe in the TSA bound form becomes faster accompanied with increased freedom for rotational motion, after ligand binding (Figure 2, Figure S3, Table S1). This unexpected change in flexibility of the dansyl probe in TSA bound state could indicate an important role of an allosteric nature by the neighboring non-active site amino acids, upon ligand binding. The Cys69-dansyl probe site is about 19 Å from C9 atom of TSA. The slow rotational component also decreased in TSA bound form, (Figure 2, Table S1) implying faster global rotation and thus asserting a structural compaction of mCM after ligand binding.

**Figure 2.**
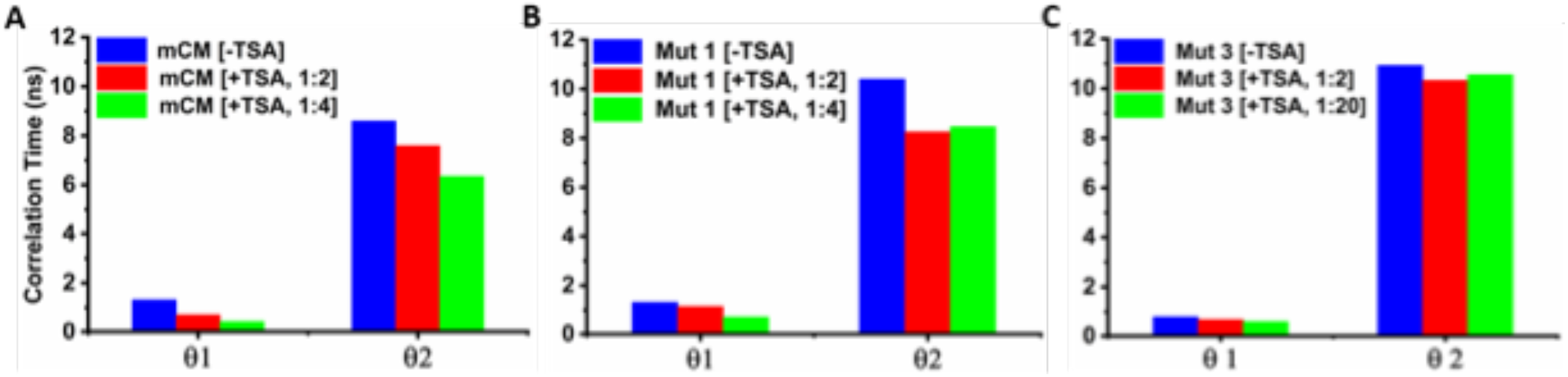
Histogram representing a comparison of the correlation time(s) in dansyl labeled mCM in its unbound and TSA bound states. (A) mCM, (B) Mutant 1 and (C) Mutant 3. The corresponding amplitude and std. dev. for each correlation time in the histogram in mentioned in Table S1.

In case of the Mutant1, a <u>fast</u> rotational component of 1.2 ns (amp=0.37) and a <u>slow</u> rotational component of 10.3 ns (amp=0.63) was obtained (Figure 2, Figure S4, Table S2). Surprisingly, Mutant1 (unbound) displayed a longer tumbling time for the slow rotational component when compared to mCM which means that mCM adopts a less compact (expanded) globular structure after undergoing mutation (W24K). This new finding from anisotropy decay data was later correlated with other structural and functional attributes of mCM to understand the impact of point mutations on mCM structure and function. After TSA binding (1:2 and 1:4 ratios), a more compact global structure of Mutant1 was indicated by the decrease in the slow rotational component to ~8.4 ns. But the compaction displayed in Mutant1 after TSA binding was less than that observed for mCM. As in mCM, ligand binding also leads to a more dynamic dansyl probe in Mutant1 (Figure 2, Table S1).

Mutant3 revealed a <u>fast</u> rotational component of 0.8 ns (amp=0.56) and a <u>slow</u> rotational component of 10.9 ns (amp = 0.44) (Figure 2, Figure S5, Table S3). It is important to note here that the dansyl probe is attached to Cys 6, a site which is far from the active site (Cys6-SH to TSA-C9 is ~12 Å) and in a different helix than Cys 69 residue. This dansyl probe (in unbound Mutant3) displayed more flexibility than in mCM and Mutant1 indicating a less hindered environment around Cys 6 residue than Cys 69 residue as evidenced by increased β_1_ (Figure S11) and faster θ_1_. Moreover, Mutant3 displayed the most expanded global structure amongst all mCM variants, inferred from its slow rotational component of ~11 ns; higher than mCM and the other mutants. After TSA binding, Mutant3 reported insignificant change in its global structure even after an excess of TSA was used (Figure 2, Table S3). Some major structural perturbations in Mutant3, owing to 3 mutations, might be the reason for its poor sensitivity to TSA. This was investigated further.

### Measuring changes in spatial proximity between Trp—Cys pair in mCM and its mutants upon ligand binding using FRET

To gather more specific information on the local structural changes in structure of mCM upon TSA binding, FRET data were acquired using time-resolved fluorescence. Analysis of the fluorescence intensity decay of the two Trp in mCM revealed the presence of three lifetimes (Figure S6). A short component of ~0.8 ns (α_1_ = 0.44), an intermediate component of ~2.2 ns (α_2_ = 0.40) and a long component of ~5.2 ns (α_3_ = 0.16). The FRET distance between Trp24/26 and Cys69 in mCM was ascertained by measuring the lifetime of the Trp residues in unlabeled versus the dansyl labeled form (Figure S6). As Trp displays multi-exponential decay, the amplitude weighted mean fluorescence lifetime was used for FRET calculations. The mean lifetime decreased from ~2 ns to ~1.3 ns. The FRET distance was calculated from these values to be ~24Å, close to the value of 23Å obtained in steady-state fluorescence experiments. The mean lifetimes in mCM after TSA binding didn’t reveal any significant change with lower concentrations of TSA, but a noticeable change in FRET distance from ~24.2 Å to ~22.3 Å was interpreted from the lifetime values when an excess of TSA (1:20 ratio) was used (Figure S6). The existence of two closely placed Trp does complicate the interpretation of fluorescence lifetime as contributions from individual Trp residues cannot be segregated.

The single Trp mutant (W24K), referred to as Mutant1 in our studies, provided a much simpler and more reliable system to monitor the structural changes in mCM in the presence of the TSA. Fluorescence decay intensity analysis of the single Trp residue presented two distinct lifetimes and a mean lifetime of ~3.4 ns (Figures 3A, B, C and Table 1). A short lifetime of ~1.8 ns (α_1_ = 0.48) and a long lifetime of ~5 ns (α_2_ = 0.52) were extracted from the decay fitting and analysis. The addition of TSA to unlabeled Mutant1 didn’t affect the lifetime of Trp significantly, as also observed in mCM (Table S4). Time-resolved FRET data from the Trp26-Cys69 (dansyl labeled) pair of Mutant1 revealed a FRET distance of ~23 Å (Figure 3, Table 1), close to the value (~24 Å) obtained for mCM. Interestingly, the addition of the TSA to this labeled Mutant1 drastically changed the FRET distance (Figure 3, Table 1). At low TSA concentrations (1:1 and 1:2 ratios), the FRET distance was brought down from ~23 Å to ~18 Å. At slightly higher TSA concentrations (1:4), the FRET distance showed a further reduction to ~16 Å. Addition of even higher concentrations of TSA (1:20) didn’t bring any further change to the FRET distance (Figure 3D). This drastic change in FRET distance can be interpreted as a tightening or collapse of the protein structure around the TSA after its binding, as the flexible loop containing the Trp26 and the helix containing Cys69 approach each other in the bound form. Another observation is the introduction of a very short lifetime ~0.18-0.09 ns in the dansyl labeled form of Mutant1. We believe that this short lifetime component may have risen due to quenching by FRET and it shows dominance with an amplitude of ~0.7-0.8. The amplitude of this very short lifetime component becomes even more dominant in the TSA bound forms.

**Table 1.** Tryptophan lifetime values (ns) for unlabeled (no TSA) and dansyl labeled (at Cys 69) Mutant1 in PBS buffer pH 7.4 with varying TSA concentrations. The calculated FRET distances are reported.

| Mutant 1 | $\tau_1^a$<br>(ns) | $\alpha_1^b$ | $\tau_2^a$<br>(ns) | $\alpha_2^b$ | $\tau_m^c$<br>(ns) | Avg. $\tau_m^d$<br>(ns) | $\chi^2^e$ |
| --- | --- | --- | --- | --- | --- | --- | --- |
| unlabeled | 1.78 | 0.48 | 4.97 | 0.52 | 3.4 | 3.4±0.1 | 1.05 |
| labeled<br>(dansyl) | 1.44 | 0.78 | 4.41 | 0.22 | 2.07 | 2.0±0.08 | 1.1 |

| Mutant1 | $\tau_1^a$<br>(ns) | $\alpha_1^b$ | $\tau_2^a$<br>(ns) | $\alpha_2^b$ | $\tau_3^a$<br>(ns) | $\alpha_3^b$ | $\tau_m^c$<br>(ns) | Avg. $\tau_m^d$<br>(ns) | $\chi^2^e$ | FRET<br>distance<br>(Å) | FRET<br>Efficiency |
| --- | --- | --- | --- | --- | --- | --- | --- | --- | --- | --- | --- |
| No TSA | 1.44 | 0.78 | 4.41 | 0.22 | --- | --- | 2.07 | 2.0 ± 0.08 | 1.1 | 23 ± 0.2 | 36.7% |
| 1:1 TSA | 1.75 | 0.15 | 5.22 | 0.09 | 0.17 | 0.76 | 0.85 | 0.85 ± 0.12 | 1.1 | 18 ± 0.3 | 71.9% |
| 1:2 TSA | 1.5 | 0.18 | 4.94 | 0.11 | 0.18 | 0.71 | 0.93 | 0.94 ± 0.04 | 1.1 | 18 ± 0.2 | 71.9% |
| 1:4 TSA | 1.62 | 0.12 | 5.1 | 0.04 | 0.11 | 0.84 | 0.49 | 0.49 ± 0.05 | 1.1 | 16 ± 0.5 | 84.0% |
| 1:20 TSA | 1.61 | 0.09 | 5.04 | 0.04 | 0.09 | 0.87 | 0.42 | 0.45 ± 0.06 | 1.1 | 16 ± 0.3 | 84.0% |

<sup>a</sup>fluorescence lifetime(s); <sup>b</sup>amplitude(s); <sup>c</sup>mean lifetime(s); <sup>d</sup>average mean lifetime(s); <sup>e</sup>reduced $\chi^2$ for the fit.

**Figure 3.**
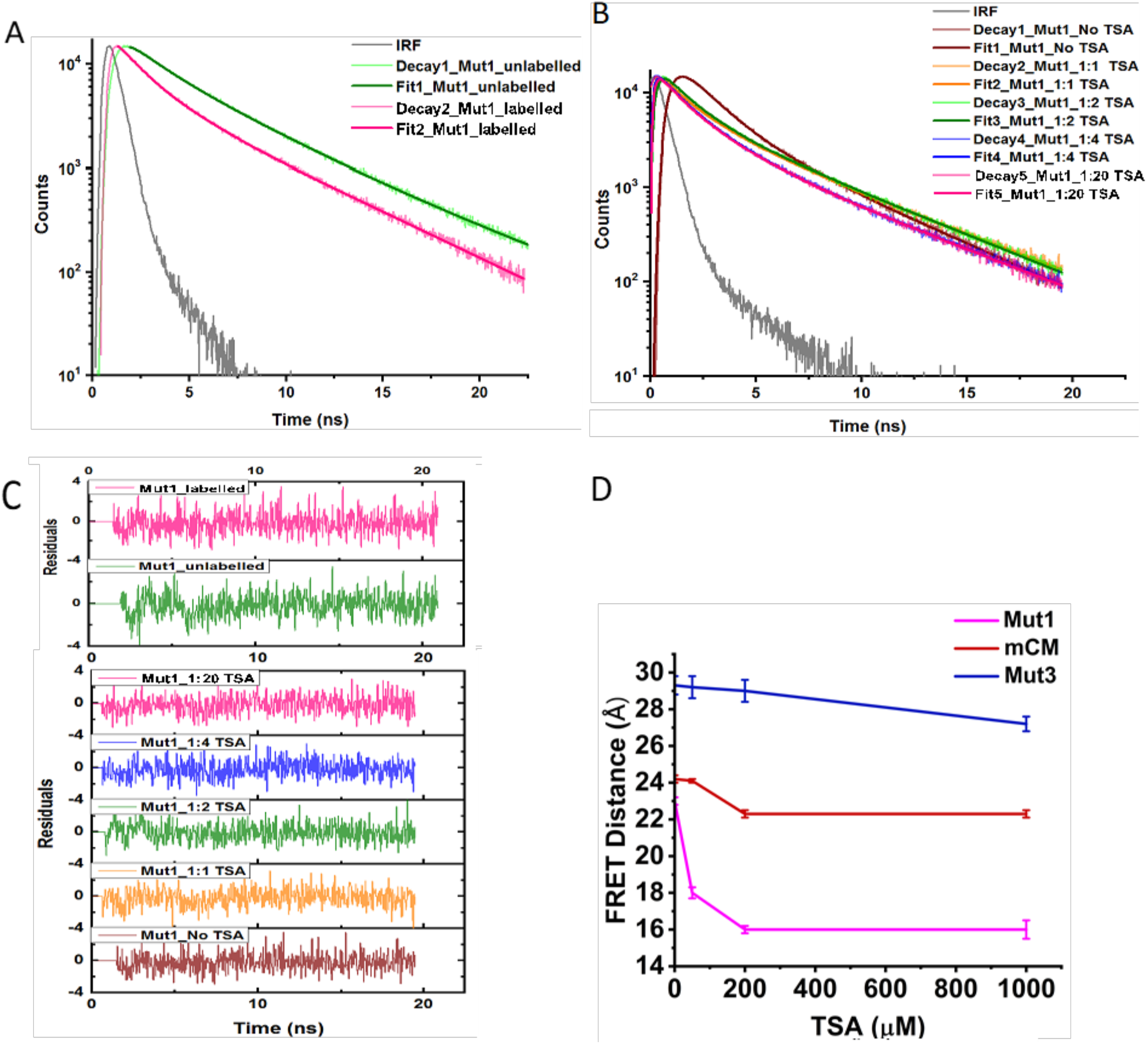
Fitted time-resolved fluorescence intensity decay profile of 50 μM dansyl labeled Mutant1 [A] unlabeled and labeled forms, [B] with varying concentrations of the TSA (50 μM, 100 μM, 200 μM and 1mM corresponding to 1:1, 1:2, 1:4 and 1:20 ratios respectively), [C] residuals for the fit and [D] FRET distance comparison in mCM, Mutant1 and Mutant3 observed before and after binding of the ligand TSA (varying concentrations).

In the triple mutant, Mutant3 (W24K/C69A/A6C), the FRET distance between Trp26-Cys6 was found to be ~29.3 Å, a distance longer than that reported for Trp26-Cys69 (Figure S8 A,D). In TSA binding experiments, high concentrations of the ligand (a 40-fold excess) were required to showcase a noticeable change in FRET distance of Trp26-Cys6 from ~29.3 Å to ~25.7 Å (Figure S8 B,D). It was intriguing to find such contrast in TSA concentrations required to bring about ligand-bound compaction in Mutant3, when compared to Mutant1. Figure 3D summarizes the effect of TSA concentration on the measured FRET distance for mCM, Mutant1 and Mutant3. It reveals that while Mutant1 is most sensitive to TSA binding, Mutant3 is least sensitive, consistent with rotational correlation data. Interestingly, mCM which has two Trp donors for FRET, reveals only a modest sensitivity, perhaps due to averaging of contribution from two Trps that may have different orientations. These results prompted us to investigate the enzyme kinetic parameters of mCM and its mutants to obtain insights on consequences of point mutations on the catalytic activity of mCM.

The double mutant, Mutant2 (W24K/C69A), does not possess a FRET pair due to absence of any Cys residue. Its Trp fluorescence lifetime in absence and presence of TSA were comparable; with average lifetime values of 3.35 ns and 3.40 ns respectively (Figure S7, Table S4).

MEM distributions of Trp fluorescence lifetime data for mCM as well as the mutants obtained by non-linear least square analysis are shown in Figure. S9. The Figure S9A shows that mCM displays only a slight shift in the peak lifetime(s) with addition of TSA. Higher amplitudes for the lifetime peaks are seen without TSA. Mutant1 in Figure S9B reveals the large presence of short lifetime (~0.1 ns) with addition of TSA specifically in the 1:4 and 1:20 ratios. In contrast, Mutant2 MEM profiles (Figure S9C) are insensitive to addition of TSA. In Mutant3 appearance of a short lifetime component is evident at higher TSA concentrations like in Mutant1. MEM lifetime distributions reveal subtle shifts in the lifetime distribution with addition of TSA which are not discernible from discrete analysis shown in Table S4.

### Enzyme activity and circular dichroism data confirm changes in the function and structure of mCM mutants, respectively

The spectrophotometric data from enzyme assays of mCM were extrapolated to obtain K_m_ (165 ± 46 μM) and k_cat_ (7.7 ± 0.17 s^−1^) values which were comparable to those reported earlier (K_m_ 180 μM, k_cat_ 3.2 s^−1^) [17]. Mutant1 gave slightly different kinetic parameters (Table 2) compared to mCM, which was unexpected, as earlier reports mention that Trp 24 to Lys 24 mutation did not impact mCM functionality (17). Some major changes in secondary structure were also observed in our CD measurements (Figure 4B), but the structure and function of mCM remained mostly conserved in Mutant1. However, Mutant2 and Mutant3 showed drastic changes in their enzyme catalytic ability. A decline of 97% and 89% in enzyme activity was observed for Mutant2 and Mutant3, respectively (Table 2). Similar deviations from mCM were also interpreted from fluorescence lifetime measurements for Mutant3, as stated earlier. Interestingly, Mutant3 had kinetic parameters closer to mCM even if it possessed one extra point mutation apart from the ones that overlapped with Mutant2. Both enzyme assays and CD showed some recovery in enzyme activity and α-helicity in Mutant3 when compared with the cysteine-less Mutant2 (Figure 4A, 4B). The introduction of cysteine at a new position (Ala6 to Cys6) in Mutant3 seems to partially compensate for the loss in structural/functional integrity. We *hypothesize* that the single cysteine residue (Cys 69) in mCM plays an important role in maintaining its structure and function. Analysis of the CD spectra by structural prediction software (K2D3) quantified the decrease in alpha-helicity from 82% in mCM to 76% in Mutant1, 54% in Mutant2 and 59% in Mutant3. A distortion in helix3 due to the cysteine to alanine mutation could be the prime cause of disruption in mCM features rather than a direct involvement of the cysteine residue with the active site or the ligand. TSA binding to mCM did not bring about any changes to the secondary structural features in mCM or its mutants (Figure S10). This suggests that effects of TSA binding are too subtle to be picked by CD.

**Table 2.**
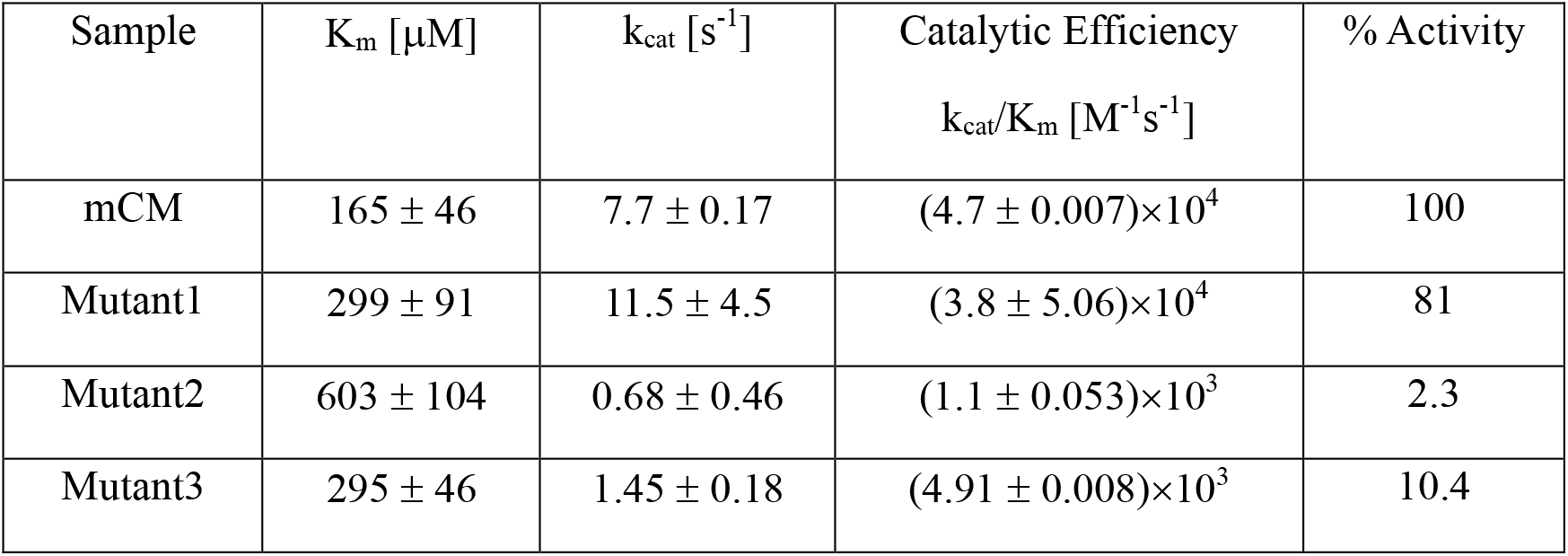
Comparison of the kinetic parameters of mCM with its mutants.

**Figure 4.**
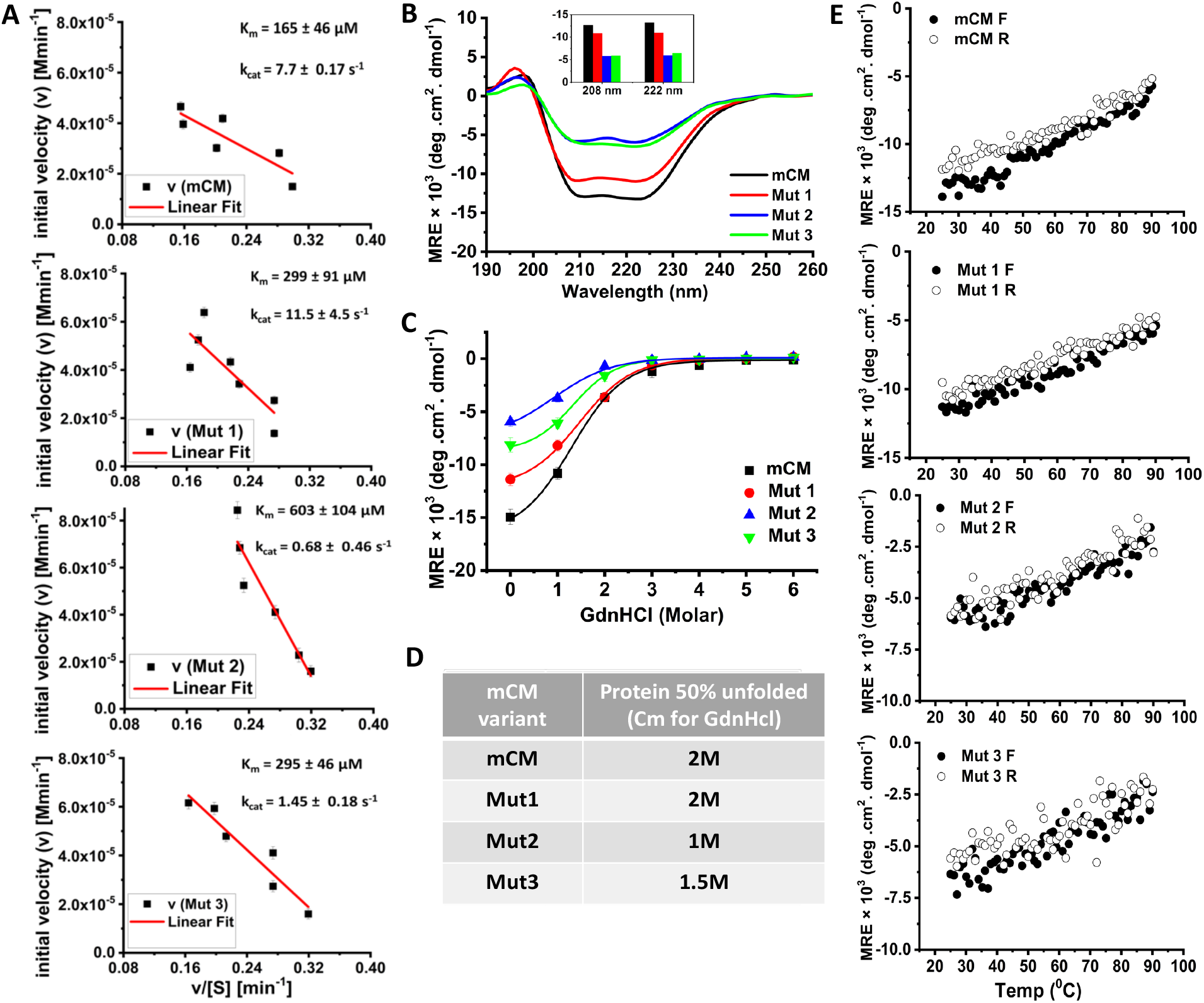
Functional and structural studies in mCM using enzyme assays and CD measurements. (A) Enzyme kinetics extrapolated as Eadie-Hofstee plot for mCM, Mutant1, Mutant2 and Mutant3 with a linear fit (red) giving the values for the slope (-K_m_), y-intercept (V_max_) and R-square. (B) CD spectra of mCM and its mutants. (Inset) The values of the two characteristics peaks of α-helices in CD spectra are compared across mCM and its mutants. (C) Chemical denaturation of mCM and its mutants monitored by UVCD at 222 nm wavelength with increasing concentrations of GdnHCl (0-6 M). Solid fitted lines represent a dose-response curve of non-linear regression analysis using OriginPro 8 software. (D) Table showing mid concentration (C_m_) for GdnHCl (0-6 M) treated protein where the protein fraction unfolded is 50% for mCM and mutants (E) Thermal melting curve of mCM following changes in the ellipticity at 222 nm wavelength. Ellipticity was monitored continuously in the increasing temperature range from 25 to 90 °C (•) and during a reverse scan from 90 to 25 °C (∘).

The unfolding curve obtained from the chemical denaturation of mCM revealed a two-state like transition, but the denatured state is achieved at different GdnHCl concentrations in all the variants (Figure 4C, 4D). mCM and Mutant1 seem to have similar stability under these denaturing conditions and they both completely lose their secondary structure beyond 4M GdnHCl. This loss in structure is observed earlier at 3M GdnHCl for Mutant3 and still earlier at 2M GdnHCl for Mutant2. A sigmoidal fit (Figure 4C) of the datasets suggests that the concentration (Cm) where the protein fraction is 50% unfolded is approximately 2 M for mCM and Mutant1, 1.5 M for Mutant3 and 1 M for Mutant2. Here, a correlation can be drawn with the secondary structural changes observed earlier in our CD spectra for mCM variants without any denaturants. An earlier report [29] showed that mCM underwent a cooperative transition from folded to unfolded state during urea-induced denaturation with the data fitting to a two-state transition model, like what we observe here. For thermal denaturation, a similar degree of reversibility was observed for mCM and all mutants (Figure 4E). Interestingly, they all appear to be noncooperative and follow a linear transition throughout the temperature scan window. A similar noncooperative thermal denaturation was previously reported for mCM (mMjCM), in the absence of any ligand [17]. They observed modest cooperativity in the TSA-bound state whereas the ordered homodimer MjCM enzyme showed a high degree of cooperativity during thermal denaturation. This lack of cooperativity in mCM in its free monomeric form is argued to be an attribute of its disordered nature.

## DISCUSSION

Intrinsically disordered enzymes, such as mCM, present a rare and compelling case to improve our understanding of how enzymes function; especially when disordered segments are involved. We set out to investigate the structural plasticity that modulates catalysis in the molten globule mCM. Our model enzyme was amenable to the sensitive fluorescence spectroscopy tools at our disposal. As per our knowledge, our report is the first to quantify the ensemble-averaged distances between chosen points in the disordered state of CM as well as the degree of change in this distance after ligand binding and structural compaction. Our time-resolved anisotropy studies supported the concept of a global collapse (ligand-induced global compaction) of the mCM structure after ligand binding (Figure 2) [30,31]. The degree of this collapse was also captured through FRET experiments (Figure 3D). We also probed the effects of rationally designed mutations namely Mutant1 (W24K), Mutant2 (W24K/C69A) and Mutant3 (W24K/C69A/A6C) on segmental dynamics, conformational adaptation, disorder-to-order transition and catalytic efficiency (Figure 4 and Table 2).

We observed that the cysteine mutants (Mutant2 and Mutant3) possessed a less compact global tertiary conformation even after ligand binding. The specific mutations in these two mutants impaired the ability of mCM to undergo effective ligand-induced disorder-to-order transitions, resulting in a decrease in global structural ordering when compared to mCM. This probably is the reason for the drastic decline of catalytic activity in these mutants. Mutant 1, with only a W24K mutation (inside a loop region) suffered minor structural changes and could function similar to mCM. A significant loss in enzyme activity and change in secondary structure in the non-active site cysteine mutants strongly suggests a role of the single cysteine (Cys69) in maintaining the secondary structure and functional integrity of the mCM enzyme. Interestingly, ANS fluorescence assay showed that the ligand can bind to the core hydrophobic region and displace ANS even in the cysteine mutants, Mutant2 and Mutant3. Thus, the cysteine mutations might not have affected ligand binding to a considerable extent. Instead, our results indicate that the hydrophobic collapse of the entire mCM structure following ligand binding might be the process severely affected by these mutations.

The free single cysteine residue (C69, helix 3) emerges as a key factor driving ligand-associated hydrophobic collapse of mCM. It is important to note that in mCM the native structure itself is affected by the C69A mutation, clearly indicated by the loss of alpha-helicity and decrease in global compaction in our studies. The re-introduction of cysteine at position 6 (A6C) in Mutant3 does not restore the structure or function of mCM to any useful extent. This clearly suggests that not only the specific amino acid (cysteine), but its correct positioning is also crucial in maintaining the structural and functional integrity of mCM. A recent *in silico* study by Mahto et al. [36] showed that in TSA bound state of mCM, a kink is formed in helix3, as a result of hydrogen bonding between D54 and TSA. This kink in turn disrupts multiple hydrogen bonds between R64 and helix1 that exists in apo mCM, in TSA bound mCM, Strikingly, while this kink did not influence the discrete analysis of Trp fluorescence intensity decay (Table S4) or CD data (Fig. S10) in TSA bound mCM, it showed subtle shifts in MEM analysis (Fig. S9A). The C69A mutation located in the vicinity of these residues in helix3, could possibly alter the structure near this kink, thereby changing local dynamics and perturbing TSA binding in the mutants. Their study also highlighted the consolidation of mCM structure beyond the active site.

An earlier study investigating the effects of point mutations at the active site of mCM [32] reported that this disordered enzyme is relatively more resilient to mutation when compared to analogous mutation of the natural thermostable ordered CM. The activity difference between the two enzymes stemmed from context-dependent, subtle changes in fine structure of active site rather than loss of secondary structural integrity or thermodynamic stability. However, in this study, a non-active site mutation showed dramatic effects in enzyme activity due to changes in secondary structure, ligand-induced global compaction and chemical/thermal denaturation states. This is intriguing as it appears that perturbation of single residues at non-active sites can have more dramatic effects than those done at the active site. This is quite in contrast to what one would anticipate of the consequences of mutations at active versus non-active sites. Another study by Lassila et al. [33] with the analogue globular *E.col*_*i*_ chorismate mutase showed that secondary active-site residues which do not directly hydrogen bond with the substrate can tolerate various single mutations without much alteration in its stability and activity. In fact, in some mutants a decrease in catalytic activity led to increased enzyme thermal stability in a stability-activity trade-off, also reported in other mutated enzymes. However, in mCM, structural stability and catalytic activity demonstrate a strong positive correlation in all the studied mutants. The structural plasticity and flexibility associated with intrinsic disorder does not shield mCM from dramatic functional impairment originating from a single mutation introduced at a site far from the active-site of the enzyme.

Current literature has limited information on the role of disordered regions and distal non-active site residues on enzyme function. One rare but well-studied case is that of the G121 mutation (*~*19 Å from active site) in DHFR, an ordered enzyme that catalyzes the reduction of dihydrofolate to tetrahydrofolate. Many reports suggest that contact pathways created by a network of interacting residues is disrupted by this remote residue, which in turn affects catalysis [34,35]. In mCM, the single C69 residue could be involved in such a network of interacting residues that promote certain protein conformation and dynamics for optimal function. The side chain thiol group of C69 may have facilitated this interaction with other residues through hydrogen bonding. As the cysteine mutants of mCM have alanine as the replacement residue, there is no side chain to facilitate such interactions, leading to the disruption of any contact network and hence its function.

The schematic in Figure 5 summarizes the structural transitions in mCM based on our experimental observations. An illustration of how the mutations and ligand binding affect mCM structure is presented in a simplified way using cylinders and linking threads. Rows (left to right) highlight the topological changes brought about by the mutations; represented as “loosened” structures. The columns (top to bottom) display the effects of ligand binding; represented as “compact” structures but with varying degree of compaction depending on the mutation status of mCM.

**Figure 5.**
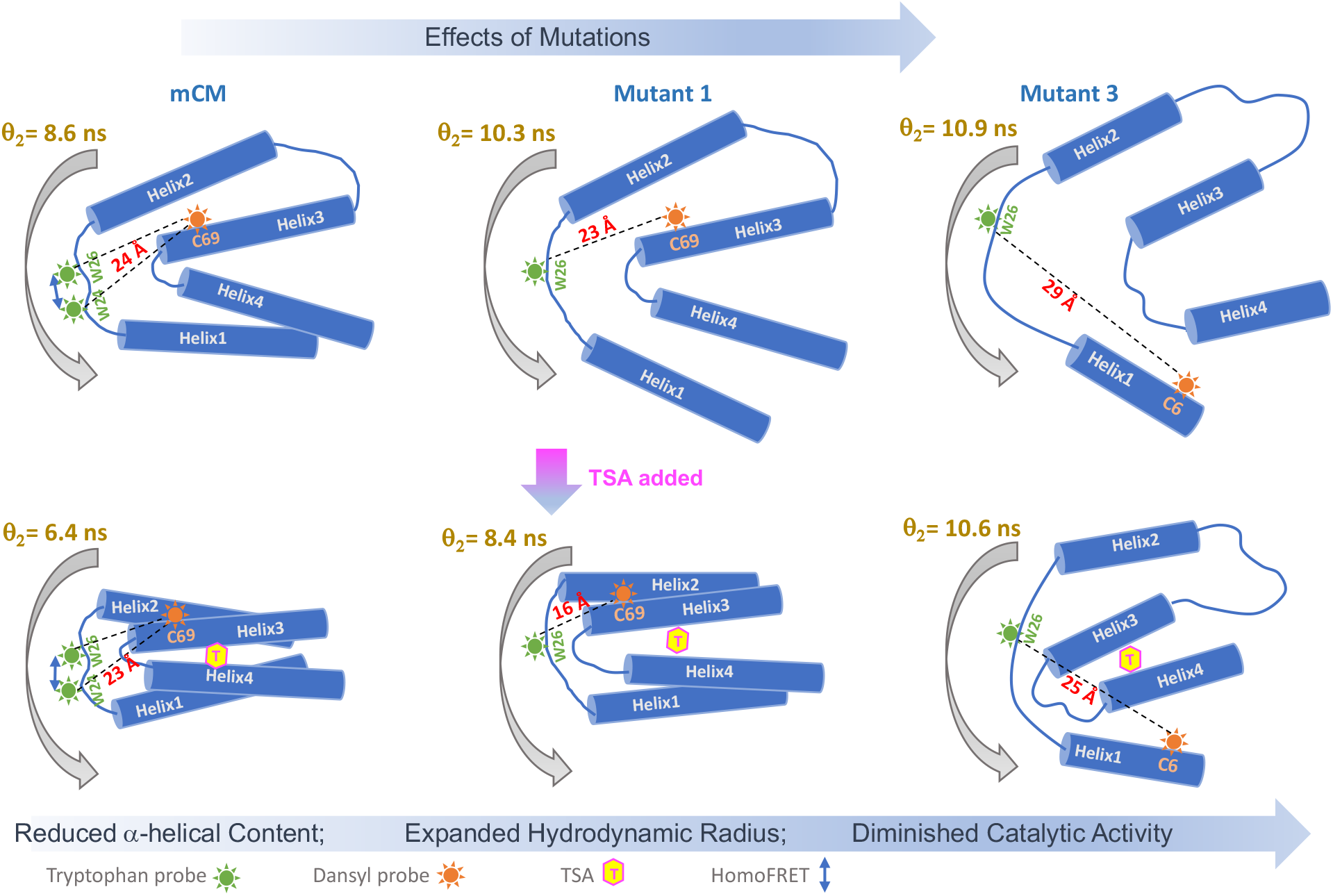
A schematic illustrating mCM transitions and the effect of mutations.

In this study, we have gathered crucial information about a highly flexible yet efficient enzyme by detailed investigations into the functional transitions made in presence of its ligand and the effects of minor perturbation on this system. The sensitivity of this enzyme to certain point mutations, located far from the active site, is one of the key highlights of this work. We believe that this can contribute in expanding the current knowledge base on this rare category of enzymes (disordered) and aid in designing better functionally viable enzymes for various pharmaceutical and industrial applications.

## CONCLUSIONS

Our investigations with mCM provided us with new insights on the structure-function relationship in a rare class of intrinsically disordered proteins that are also functional enzymes. We report on the extent of structural change in mCM and its mutants, upon binding of transition state analogue, by measuring inter-segmental motion and global compaction using time-resolved measurements of FRET-induced Trp quenching and fluorescence anisotropy of Cys-conjugated dansyl probe. The changes in structural dynamics and the resulting alterations in disorder to order transitions in all mutants of mCM could be correlated to their diminishing enzyme catalytic efficiencies. Through a detailed comparison of mCM and its mutants using FRET, fluorescence anisotropy, CD and enzyme kinetics, the drastic changes observed in mCM could be pinpointed to the single amino acid mutation of Cys 69 to Ala 69, shedding light on the crucial role played by the single cysteine residue in mCM. It was intriguing to discover that a single non-active site mutation can significantly affect structure and function in a molten globule IDP; a class of proteins known for their high tolerance to amino acid substitutions. Even more surprising is the subtle but noticeable changes in Mutant1, where the mutation (W24K) is in a highly flexible loop region. Our study highlights the susceptibility of mCM to specific mutations in non-active site locations, suggesting the importance of long-range residue interactions and secondary structure elements in maintaining the structure and function of mCM.

## ACKNOWLEDGEMENTS

The authors thank the Department of Biotechnology, Government of India, for funding this project (Ref: BT/409/NE/U-Excel/2013). The authors sincerely appreciate Prof. Donald Hilvert (ETH Zurich) for providing the mMjCM plasmid as a gift. The authors are very grateful to Prof. S.S. Ghosh (IIT Guwahati) for access to the Circular Dichroism (CD) facility at the Center of Excellence, Department of BSBE and Central Instruments Facility (CIF), IIT Guwahati, for the Mass Spectroscopy (MALDI-TOF) facility. The authors are deeply indebted to Prof. N. Periasamy (retired TIFR Mumbai) for the fluorescence intensity decay analysis software.

**Electronic Supplementary Information is available**.

## Notes

### Competing Interest Statement

The authors have declared no competing interest.

