## Supplementary Information file for "Investigating the Consequences of Non-active site Mutations on the Structure, Function and Dynamics of the Molten Globule Enzyme Monomeric Chorismate Mutase"

and

[3]Academy of Scientific and Innovative Research (AcSIR), Ghaziabad 201002, India

| Item | Contents | Page |
|---|---|---|
| Figure S1 | Trp24—Trp26 proximity in mCM and likelihood of Homo-FRET | 3 |
| Figure S2 | Trp fluorescence emission spectra of mCM (unlabeled) with TSA. | 4 |
| Figure S3 | Dansyl fluorescence anisotropy decays with fits of mCM with TSA | 5 |
| Table S1 | Anisotropy decay fitted parameters of dansyl labeled mCM (-/+ TSA) | 5 |
| Figure S4 | Dansyl fluorescence anisotropy decays with fits of mCM Mutant1 with TSA | 6 |
| Table S2 | Anisotropy decay fitted parameters of dansyl labeled mCM Mutant1 (-/+ TSA) | 6 |
| Figure S5 | Dansyl fluorescence anisotropy decays with fits of mCM Mutant3 with TSA | 7 |
| Table S3 | Anisotropy decay fitted parameters of dansyl labeled mCM Mutant3 (-/+ TSA) | 7 |
| Figure S6 | Trp fluorescence intensity decays of mCM with(out) FRET acceptor and TSA | 8 |
| Figure S7 | Trp fluorescence intensity decays of mCM Mutant2 with(out) TSA | 9 |
| Figure S8 | Trp fluorescence intensity decays of mCM Mutant3 with(out) FRET acceptor and TSA | 10 |
| Figure S9 | MEM distributions for Trp fluorescence lifetime in mCM and mutants (-/+ TSA) | 11 |
| Figure S10 | CD spectra of mCM variants (-/+ TSA) | 12 |
| Figure S11 | Plots of beta1 values from anisotropy decay analysis for mCM variants (-/+ TSA) | 12 |
| Table S4 | Trp fluorescence lifetime values for unlabeled mCM variants | 13 |
| Synthesis of Transition State Analogue (TSA) | | 14 |

A

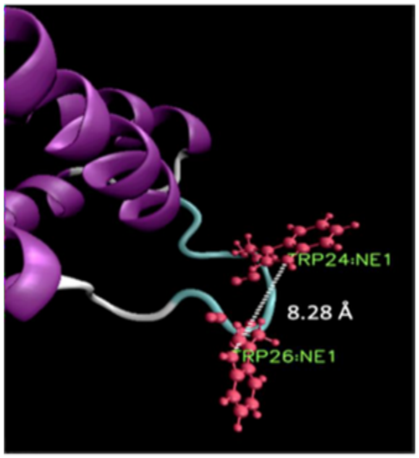


B

| Sample* | Anisotropy ($r_{ss}$) @ 295 nm | Anisotropy ($r_{ss}$) @ 310 nm | Ratio 310/295 |
|---|---|---|---|
| mCM | 0.099 ± 0.004 | 0.243 ± 0.01 | 2.42 |
| Mutant1 | 0.085 ± 0.005 | 0.137 ± 0.008 | 1.61 |
| NATA | 0.007 ± 0.001 | 0.009 ± 0.001 | 1.12 |

*10 µM sample (each) was taken in PBS buffer pH 7.2 at 25 °C for the measurements

**Figure S1.** [A] NMR structure (PDB: *2GTV*) of mCM focusing on the Trp 24 and Trp 26 residues in a ball and stick representation (red). The distance between the two tryptophan (NE1:NE1) is calculated to be approximately 8.28 Å (VMD program). [B] The table above highlights the difference in the 310/295 ratio in the two-tryptophan containing mCM and the single tryptophan variant, Mutant1. A value above 2 (2.42) recorded for mCM suggests the possibility of homoFRET between Trp 24 and Trp 26 [*Moens PD, Helms MK, Jameson DM. Detection of tryptophan to tryptophan energy transfer in proteins. Protein J. 2004 23(1):79-83. doi: 10.1023/b:jopc.0000016261.97474.2e.*]. The Förster distance ($R_0$) for Trp-Trp homotransfer is reported to be 6-12 Å [*Van Der Meer B W, Coker G and Chen S Y S 1991 Resonance energy transfer: theory and data (Wiley-VCH: New York*], suggesting that 8.3 Å separation between the indoles here, favours homoFRET. Moreover, a fairly blue-shifted emission (λmax=340 nm) in mCM favours homoFRET between Trp 24 and Trp 26 (Lakowicz, 2006).

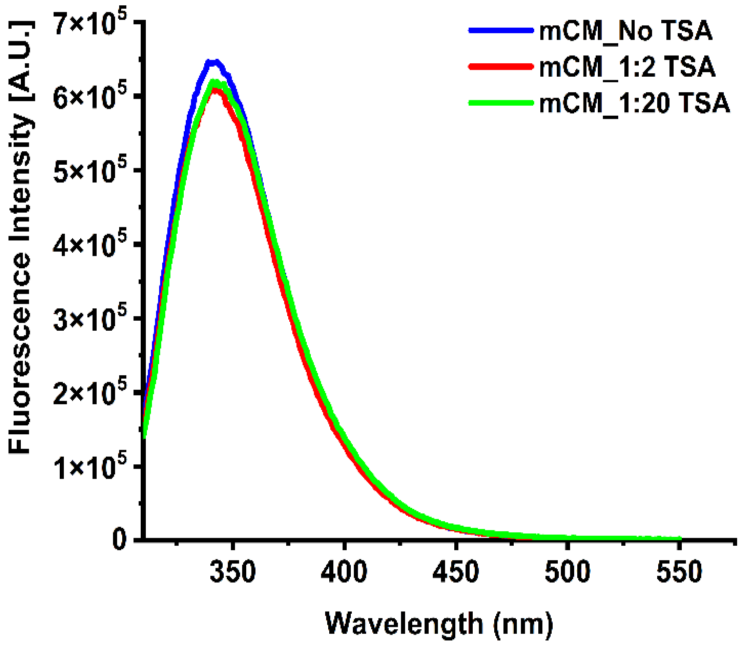


**Figure S2.** Tryptophan fluorescence spectra of mCM (unlabeled) in the presence or absence of TSA.

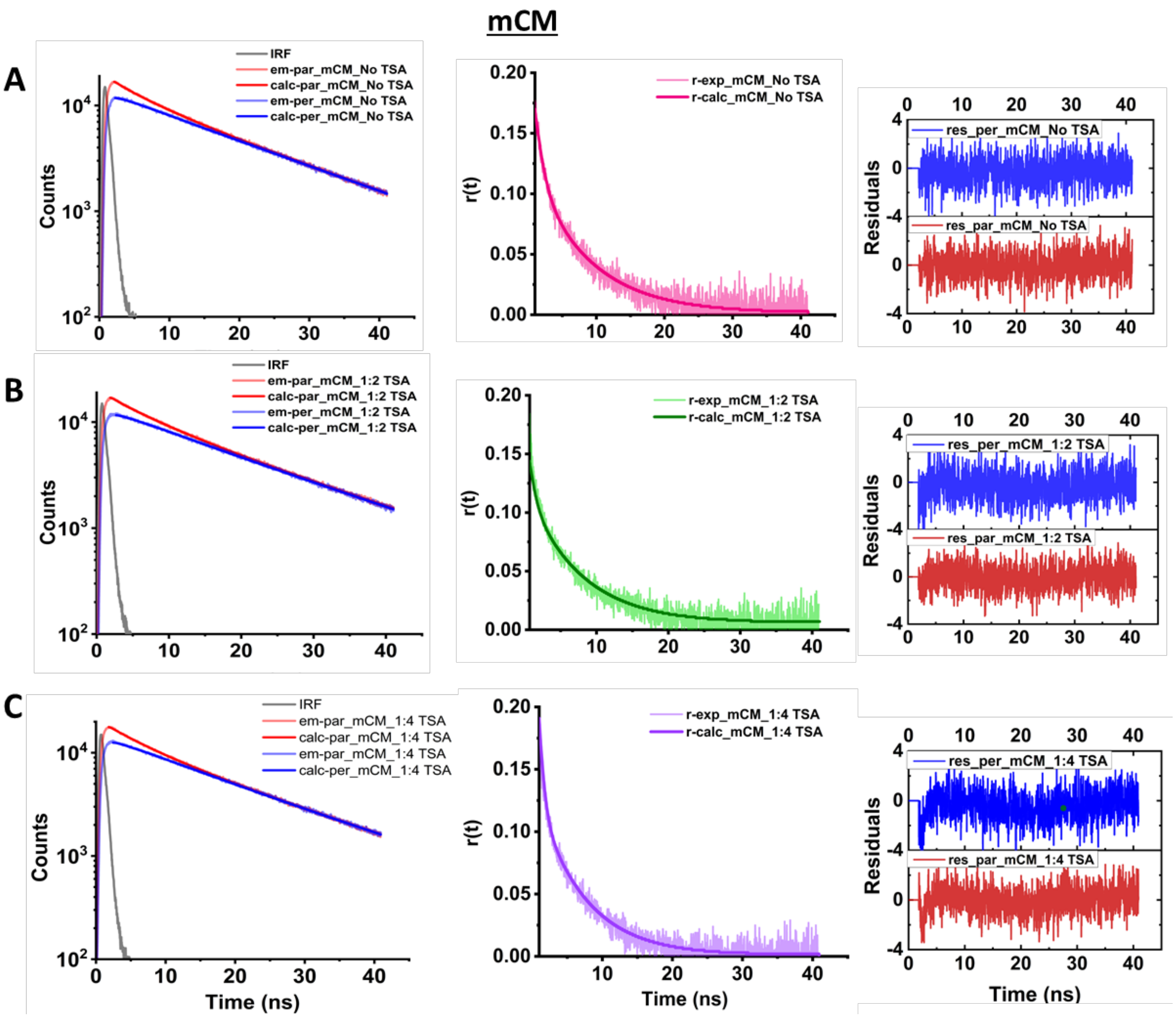


**Figure S3.** Dansyl anisotropy decay profile of mCM in unbound and bound forms (with TSA). Residuals for the fitted data are shown in the right column. Decay plots, r(t) and residuals for fitted data of dansyl probe in mCM with no added TSA (Top row, A); 1: 2 ratio TSA (Middle row, B); and 1: 4 ratio TSA (Bottom row, C) are shown.

**Table S1.** Anisotropy decay fitted parameters of dansyl labeled mCM (-/+ TSA) in PBS buffer pH 7.4

| mCM | $r_0^a$ | $\theta_1^b$ (ns) | $\beta_1^c$ | $\theta_2^b$ (ns) | $\beta_2^c$ | $\chi^{2d}$ |
|---|---|---|---|---|---|---|
| No TSA | 0.18 | 1.32 ± 0.04 | 0.44 | 8.60 ± 0.15 | 0.56 | 1.2 |
| 1:2 TSA | 0.19 | 0.71 ± 0.01 | 0.50 | 7.60 ± 0.09 | 0.50 | 1.2 |
| 1:4 TSA | 0.17 | 0.44 ± 0.01 | 0.61 | 6.35 ± 0.06 | 0.39 | 1.3 |

[a] initial anisotropy; [b] rotational correlation time(s); [c] fractional amplitude associated with the correlation time; [d] reduced $\chi^2$ for the fit

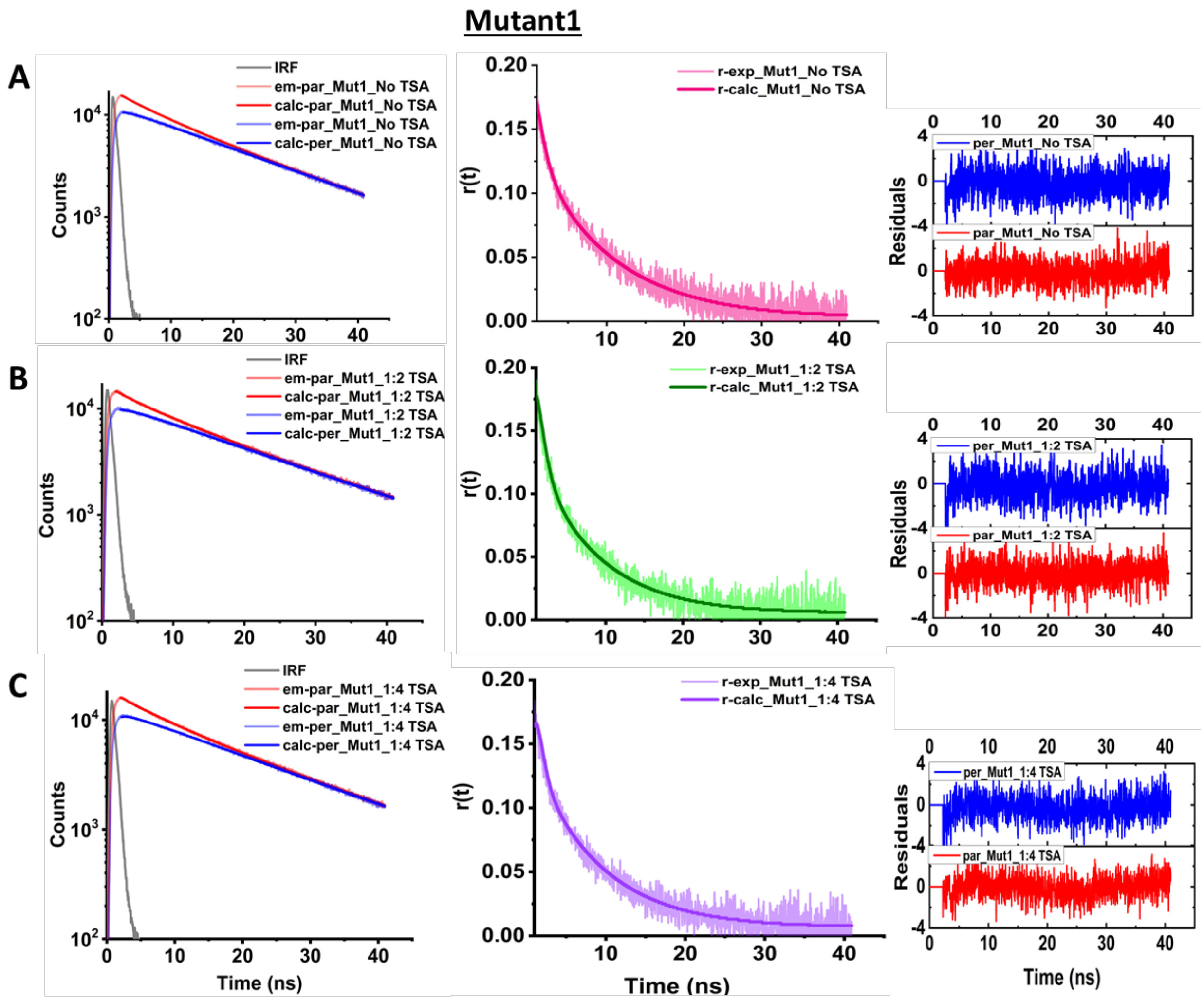


**Figure S4.** Dansyl anisotropy decay profile of Mutant1 in unbound and bound forms (with TSA). Residuals for the fitted data are shown in the right column. Decay plots, r(t) and residuals for fitted data of dansyl probe in Mutant1 with no added TSA (Top row, A); 1:2 ratio TSA (Middle row, B); and 1:20 ratio TSA (Bottom row, C) are shown.

**Table S2**. Anisotropy decay fitted parameters of dansyl labeled Mutant1 (-/+ TSA) in PBS buffer pH 7.4

| **Mutant1** | $r_0^a$ | $\theta_1^b$ (ns) | $\beta_1^c$ | $\theta_2^b$ (ns) | $\beta_2^c$ | $\chi^{2d}$ |
|---|---|---|---|---|---|---|
| No TSA | 0.18 | 1.20 ± 0.06 | 0.37 | 10.30 ± 0.15 | 0.63 | 1.17 |
| 1:2 TSA | 0.19 | 0.98 ± 0.03 | 0.49 | 8.25 ± 0.13 | 0.51 | 1.23 |
| 1:4 TSA | 0.19 | 0.72 ± 0.02 | 0.52 | 8.47 ± 0.13 | 0.48 | 1.23 |

[a] initial anisotropy; [b] rotational correlation time(s); [c] fractional amplitude associated with the correlation time; [d] reduced $\chi^2$ for the fit

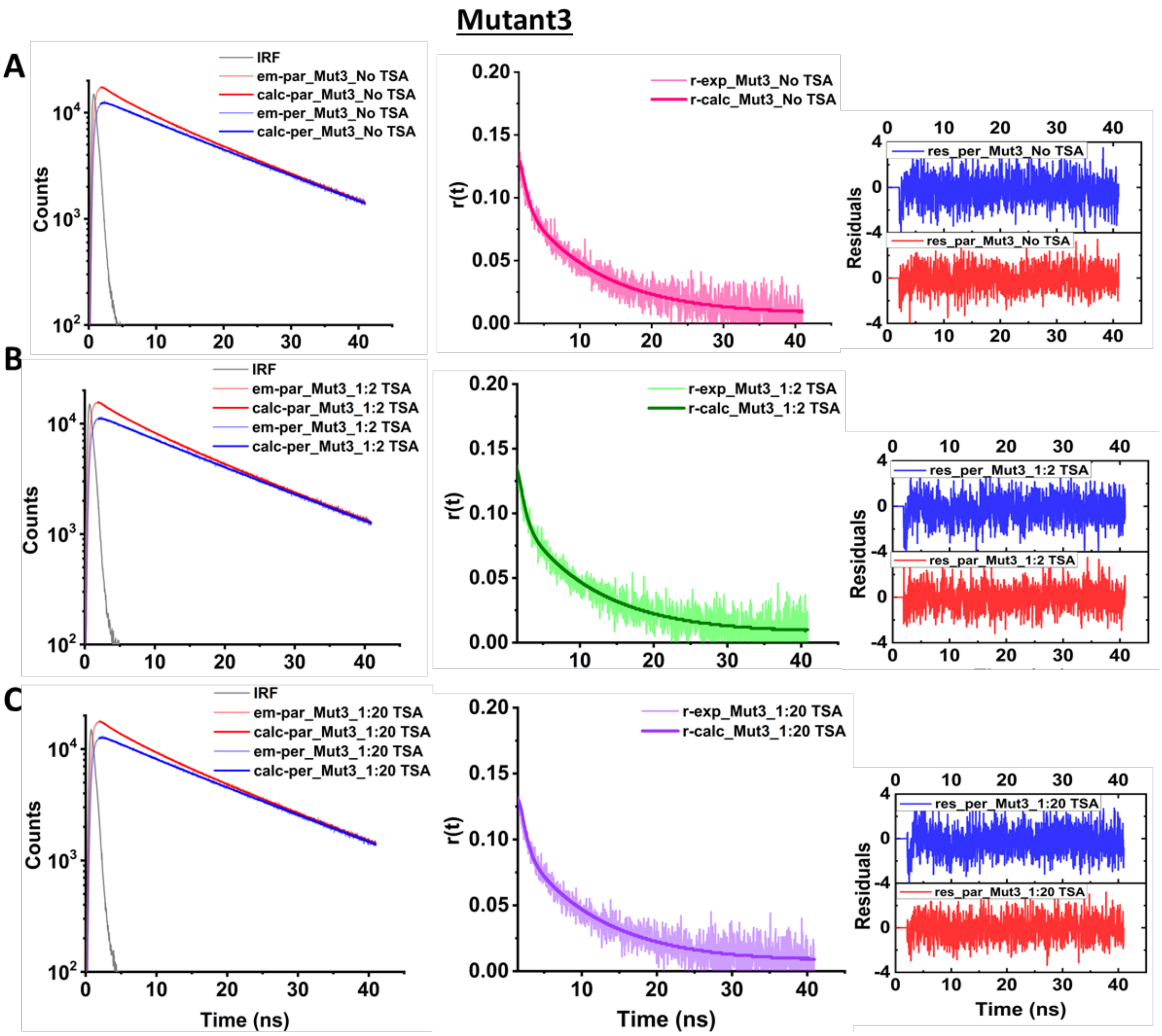


**Figure S5.** Dansyl anisotropy decay profile of Mutant3 in unbound and bound forms (with TSA). Decay plots, r(t) and residuals for fitted data of dansyl probe in Mutant3 with no added TSA (Top row, A); 1:2 ratio TSA (Middle row, B); and 1:20 ratio TSA (Bottom row, C) are shown.

**Table S3**. Anisotropy decay fitted parameters of dansyl labeled Mutant3 (-/+ TSA) in PBS buffer pH 7.4

| **Mutant3** | $r_0$[a] | $\theta_1$[b] (ns) | $\beta_1$[c] | $\theta_2$[b] (ns) | $\beta_2$[c] | $\chi^2$[d] |
|---|---|---|---|---|---|---|
| No TSA | 0.15 | 0.80 ± 0.04 | 0.56 | 10.93 ± 0.16 | 0.44 | 1.19 |
| 1:2 TSA | 0.16 | 0.70 ± 0.04 | 0.61 | 10.33 ± 0.26 | 0.39 | 1.19 |
| 1:4 TSA | 0.16 | 0.61 ± 0.03 | 0.66 | 10.56 ± 0.22 | 0.34 | 1.21 |

[a] initial anisotropy; [b] rotational correlation time(s); [c] fractional amplitude associated with the correlation time; [d] reduced $\chi^2$ for the fit

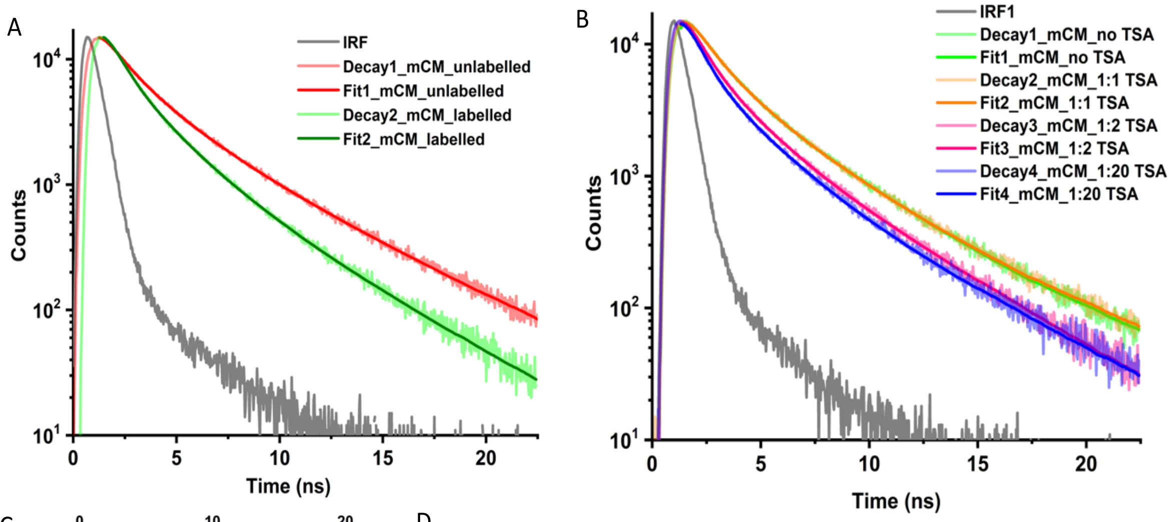


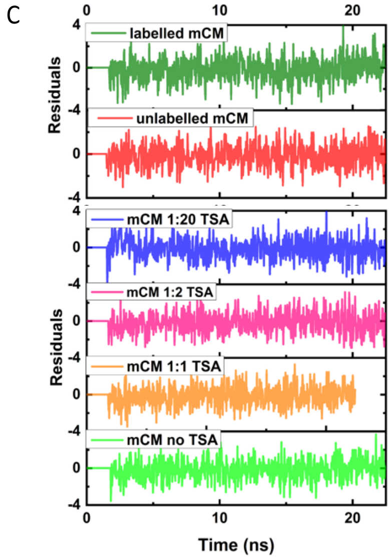


D

| mCM | $\tau_1^a$ (ns) | $\alpha_1^b$ | $\tau_2^a$ (ns) | $\alpha_2^b$ | $\tau_3^a$ (ns) | $\alpha_3^b$ | $\tau_m^c$ (ns) | Avg. $\tau_m^d$ (ns) | $\chi^{2e}$ |
|---|---|---|---|---|---|---|---|---|---|
| **Unlabeled** | 0.83 | 0.44 | 2.2 | 0.40 | 5.25 | 0.16 | 2.08 | 2.0 ± 0.2 | 1.07 |
| **Labeled (dansyl)** | 0.52 | 0.57 | 1.68 | 0.31 | 4.08 | 0.12 | 1.3 | 1.3 ± 0.1 | 1.1 |

| mCM | $\tau_1^a$ (ns) | $\alpha_1^b$ | $\tau_2^a$ (ns) | $\alpha_2^b$ | $\tau_3^a$ (ns) | $\alpha_3^b$ | $\tau_m^c$ (ns) | Avg. $\tau_m^d$ (ns) | $\chi^{2e}$ | FRET Distance (Å) |
|---|---|---|---|---|---|---|---|---|---|---|
| **No TSA** | 0.52 | 0.57 | 1.68 | 0.31 | 4.08 | 0.12 | 1.3 | 1.3 ± 0.1 | 1.1 | 24.2 ± 0.2 |
| **1:1 TSA** | 0.53 | 0.62 | 1.87 | 0.29 | 4.59 | 0.09 | 1.27 | 1.3 ± 0.04 | 1.1 | 24.1 ± 0.1 |
| **1:2 TSA** | 0.41 | 0.69 | 1.68 | 0.23 | 4.49 | 0.07 | 1.0 | 1.31 ± 0.09 | 1.1 | 22.3 ± 0.2 |
| **1:20 TSA** | 0.48 | 0.69 | 1.84 | 0.24 | 4.68 | 0.06 | 1.0 | 1.1 ± 0.1 | 1.1 | 22.3 ± 0.2 |

a: fluorescence lifetime(s); b: amptitude(s); c: mean lifetime(s); d: average mean lifetime(s);
e: reduced $\chi^2$ for the fit

**Figure S6.** Fitted Trp DONOR time-resolved fluorescence intensity decay profile of 50 μM mCM in [A] unlabeled (*donor alone*) and labelled (*donor + acceptor*) forms, [B] with varying concentrations of the TSA (50 μM, 100 μM and 1mM corresponding to 1:1, 1:2 and 1:20 ratios respectively, [C] residuals for the fit and [D] table containing Trp lifetime values (ns) for unlabeled (no TSA) and dansyl labelled (at Cys 69) mCM with varying TSA concentrations.

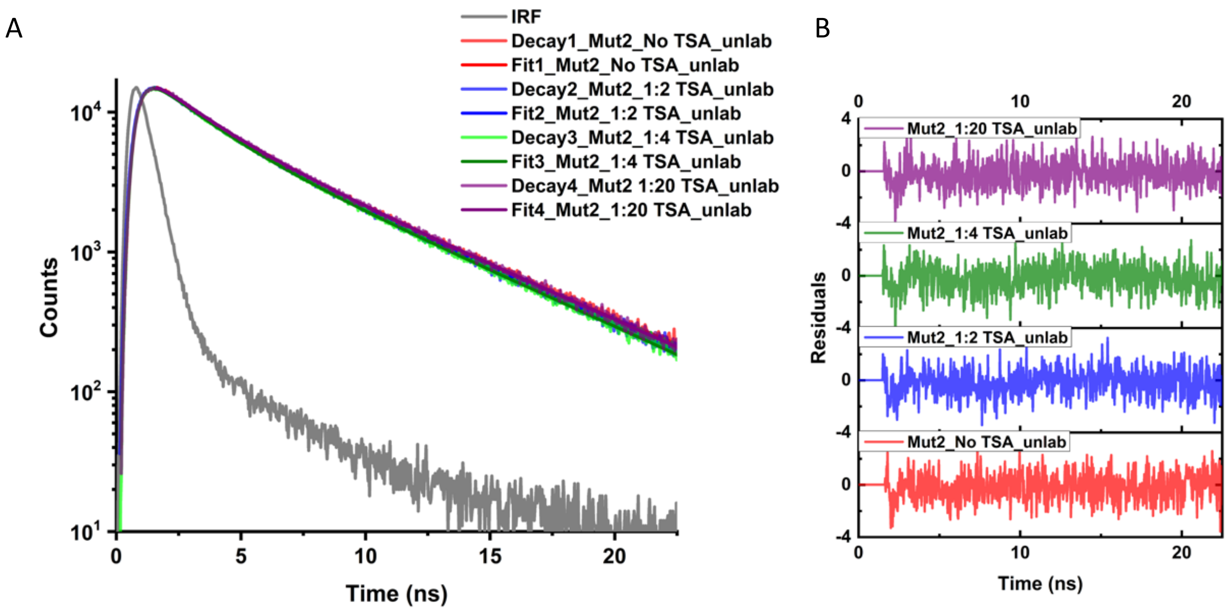


C

| Mutant2 | $\tau_1^a$(ns) | $\alpha_1^b$ | $\tau_2^a$(ns) | $\alpha_2^b$ | $\tau_m^c$ (ns) | Avg. $\tau_m^d$ (ns) | $\chi^{2e}$ |
|---|---|---|---|---|---|---|---|
| **No TSA** | 1.72 | 0.50 | 4.97 | 0.50 | 3.33 | 3.35 ± 0.04 | 1.07 |
| **1:2 TSA** | 1.90 | 0.56 | 5.25 | 0.44 | 3.38 | 3.35 ± 0.05 | 1.1 |
| **1:4 TSA** | 1.87 | 0.56 | 5.17 | 0.44 | 3.34 | 3.35 ± 0.06 | 1.06 |
| **1:20 TSA** | 1.85 | 0.52 | 5.16 | 0.48 | 3.44 | 3.40 ± 0.06 | 1.03 |

a: fluorescence lifetime(s); b: amplitude(s); c: mean lifetime(s);
d: average mean lifetime(s); e: reduced $\chi^2$ for the fit

**Figure S7.** Fitted Trp time-resolved fluorescence intensity decay profile of 50 μM Mutant2 (unlabelled) with varying concentrations of the TSA 50 μM, 100 μM, 200 μM and 1 mM corresponding to 1:1, 1:2, 1:4 and 1:20 ratios respectively [A]; residuals for the fit [B]; and a table containing Trp lifetime values (ns) [C].

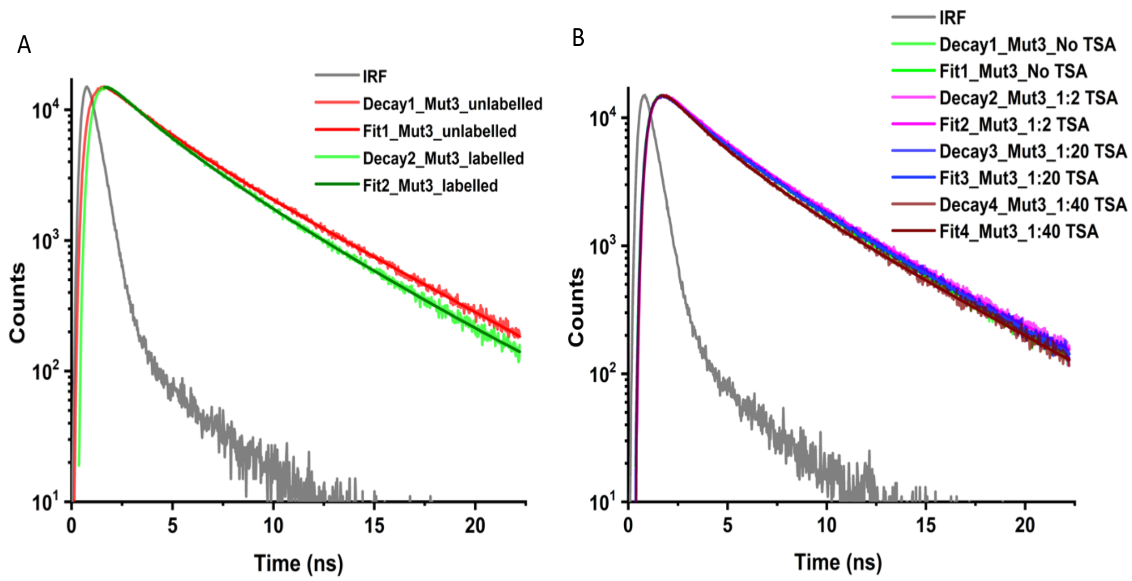


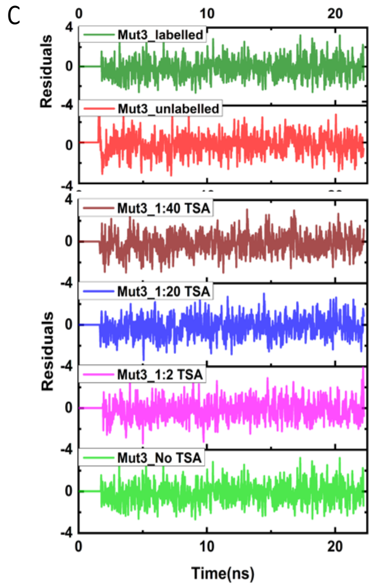


D

| Mutant3 | $\tau_1{}^a$ (ns) | $\alpha_1{}^b$ | $\tau_2{}^a$ (ns) | $\alpha_2{}^b$ | $\tau_3{}^a$ (ns) | $\alpha_3{}^b$ | $\tau_m{}^c$ (ns) | Avg. $\tau_m{}^d$ (ns) | $\chi^{2e}$ |
|---|---|---|---|---|---|---|---|---|---|
| Unlabeled | 2.15 | 0.52 | 5.20 | 0.48 | --- | --- | 3.60 | 3.5 ± 0.1 | 1.03 |
| Labeled (dansyl) | 2.92 | 0.49 | 5.47 | 0.23 | 1.10 | 0.27 | 3.03 | 3.0 ± 0.4 | 1.02 |

| Mutant3 | $\tau_1{}^a$ (ns) | $\alpha_1{}^b$ | $\tau_2{}^a$ (ns) | $\alpha_2{}^b$ | $\tau_3{}^a$ (ns) | $\alpha_3{}^b$ | $\tau_m{}^c$ (ns) | Avg. $\tau_m{}^d$ (ns) | $\chi^{2e}$ | FRET Distance (Å) |
|---|---|---|---|---|---|---|---|---|---|---|
| No TSA | 2.92 | 0.49 | 5.47 | 0.23 | 1.10 | 0.27 | 3.03 | 3.00 ± 0.40 | 1.02 | 29.3 ± 0.5 |
| 1:1 TSA | 2.52 | 0.44 | 5.23 | 0.32 | 0.85 | 0.23 | 3.01 | 3.00 ± 0.06 | 1.03 | 29.2 ± 0.6 |
| 1:20 TSA | 2.47 | 0.46 | 5.29 | 0.29 | 0.77 | 0.24 | 2.87 | 2.85 ± 0.02 | 1.04 | 27.2 ± 0.4 |
| 1:40 TSA | 2.23 | 0.46 | 5.31 | 0.25 | 0.74 | 0.24 | 2.60 | 2.60 ± 0.05 | 1.03 | 25.7 ± 0.6 |

a: fluorescence lifetime(s); b: amptitude(s); c: mean lifetime(s); d: average mean lifetime(s); e: reduced $\chi^2$ for the fit

**Figure S8.** Fitted Trp DONOR time-resolved fluorescence intensity decay profile of 50 µM Mutant3 in [A] unlabeled and labeled forms, [B] with varying concentrations of the TSA (100 µM, 1mM and 2 mM corresponding to 1:2, 1:20 and 1:40 ratios respectively, [C] residuals for the fit and [D] table containing Trp lifetime values (ns) for unlabeled (no TSA) and dansyl labeled (at Cys 6) Mutant3 with varying TSA concentrations.

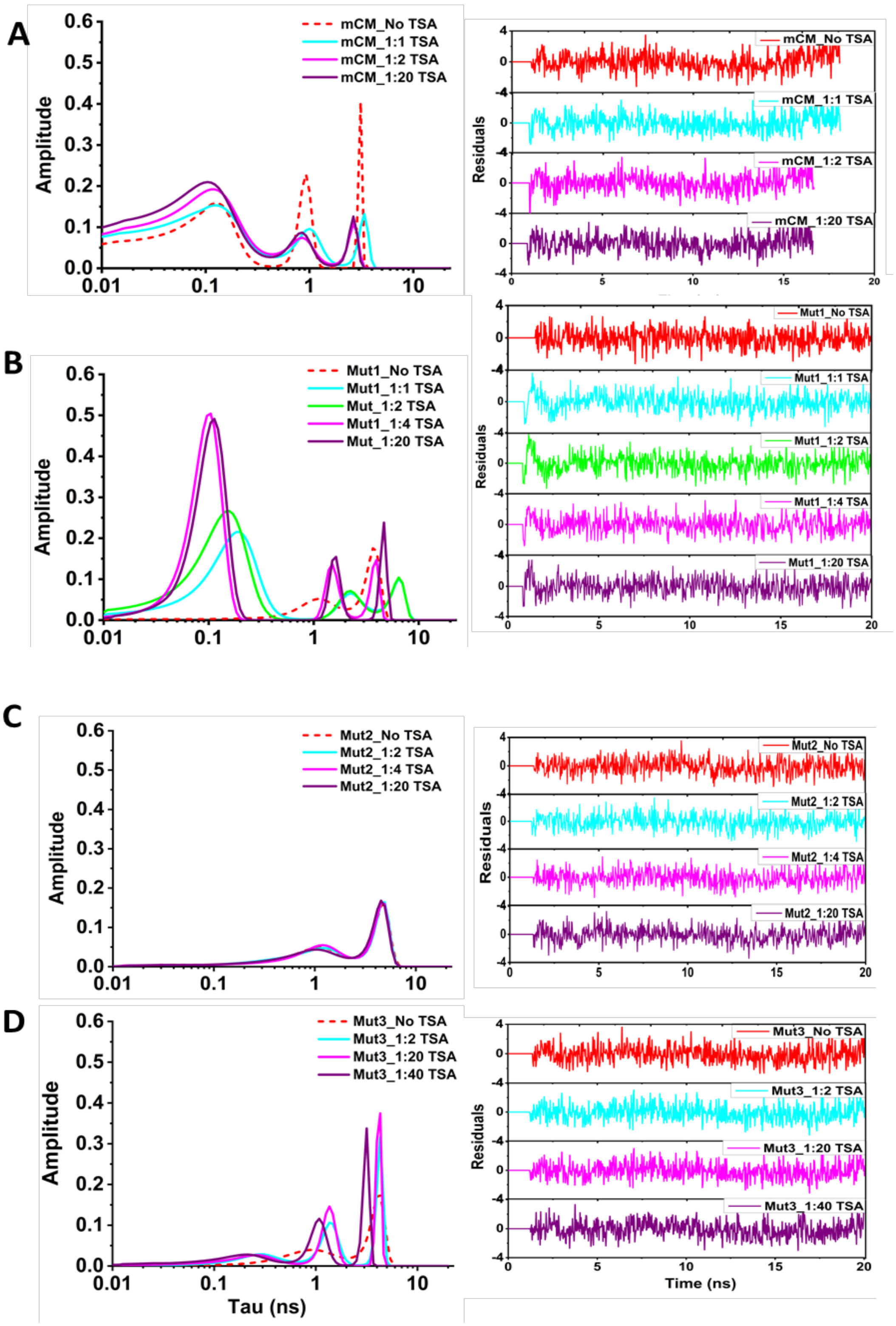


**Figure S9.** MEM distributions for Trptophan fluorescence lifetime in mCM and its mutants in the absence/presence of different protein: TSA ratios. [A] mCM; [B] Mutant1; [C] Mutant 2; [D] Mutant3;.

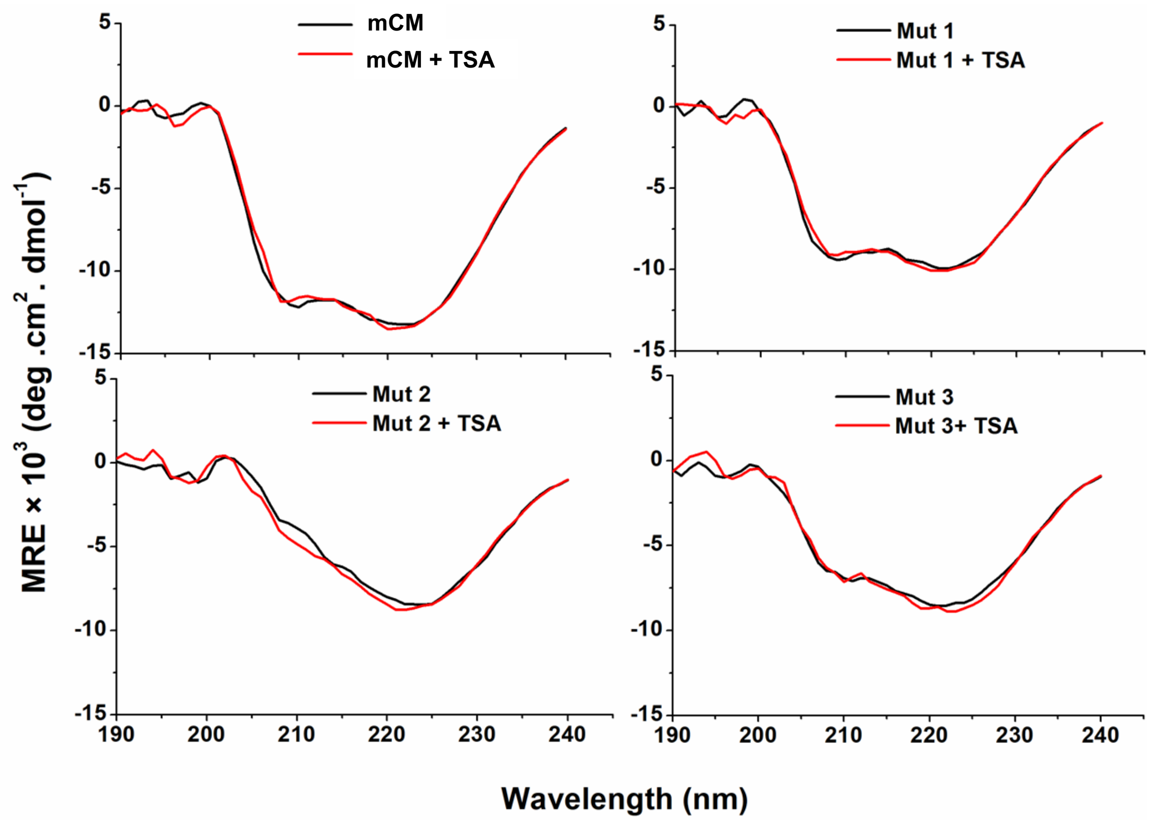


**Figure S10.** CD spectra of mCM and its variants (15 µM) in the presence and absence of TSA (60 µM).

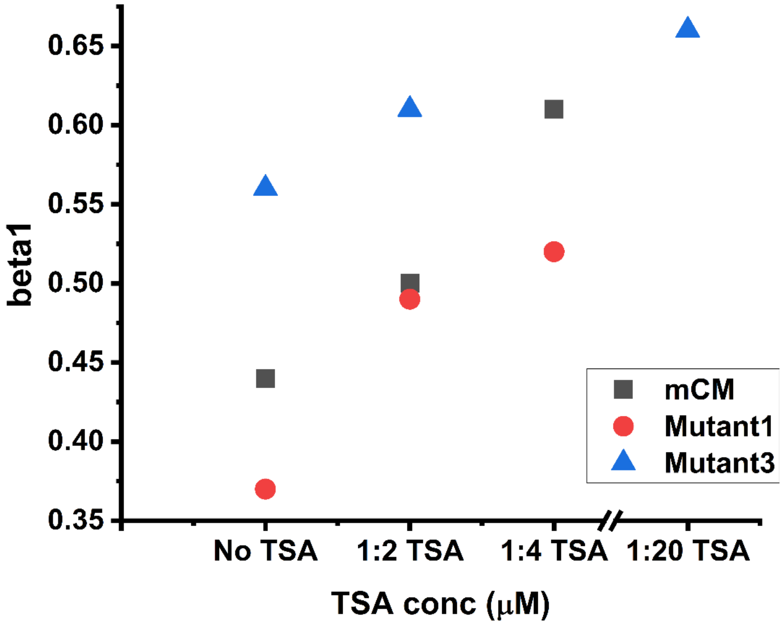


**Figure S11.** Plots of beta1 values from anisotropy decay analysis for mCM and its mutants at various TSA concentrations

**Table S4**. Tryptophan fluorescence lifetime values (ns) for unlabeled mCM and mutants (20 μM) in the presence or absence of TSA (1mM)

| mCM | $\tau_1$[a] (ns) | $\alpha_1$[b] | $\tau_2$[a] (ns) | $\alpha_2$[b] | $\tau_3$[a] (ns) | $\alpha_3$[b] | $\tau_m$[c] (ns) | Avg. $\tau_m$[d] (ns) | $\chi^2$[e] |
|---|---|---|---|---|---|---|---|---|---|
| TSA (-) | 0.83 | 0.44 | 2.2 | 0.40 | 5.25 | 0.16 | 2.08 | 2.0 ± 0.2 | 1.07 |
| TSA (+) | 0.82 | 0.45 | 2.2 | 0.40 | 5.20 | 0.17 | 2.13 | 2.1 ± 0.1 | 1.10 |

| Mutant1 | $\tau_1$[a] (ns) | $\alpha_1$[b] | $\tau_2$[a] (ns) | $\alpha_2$[b] | $\tau_3$[a] (ns) | $\alpha_3$[b] | $\tau_m$[c] (ns) | Avg. $\tau_m$[d] (ns) | $\chi^2$[e] |
|---|---|---|---|---|---|---|---|---|---|
| TSA (-) | 1.78 | 0.48 | 4.97 | 0.52 | --- | --- | 3.40 | 3.4 ± 0.1 | 1.05 |
| TSA (+) | 1.80 | 0.46 | 5.06 | 0.54 | --- | --- | 3.56 | 3.5 ± 0.1 | 1.10 |

| Mutant2 | $\tau_1$[a] (ns) | $\alpha_1$[b] | $\tau_2$[a] (ns) | $\alpha_2$[b] | $\tau_3$[a] (ns) | $\alpha_3$[b] | $\tau_m$[c] (ns) | Avg. $\tau_m$[d] (ns) | $\chi^2$[e] |
|---|---|---|---|---|---|---|---|---|---|
| TSA (-) | 1.72 | 0.50 | 4.97 | 0.50 | --- | --- | 3.33 | 3.35 ± 0.04 | 1.05 |
| TSA (+) | 1.85 | 0.52 | 5.16 | 0.48 | --- | --- | 3.44 | 3.40 ± 0.08 | 1.03 |

| Mutant3 | $\tau_1$[a] (ns) | $\alpha_1$[b] | $\tau_2$[a] (ns) | $\alpha_2$[b] | $\tau_3$[a] (ns) | $\alpha_3$[b] | $\tau_m$[c] (ns) | Avg. $\tau_m$[d] (ns) | $\chi^2$[e] |
|---|---|---|---|---|---|---|---|---|---|
| TSA (-) | 2.15 | 0.52 | 5.20 | 0.48 | --- | --- | 3.60 | 3.50 ± 0.15 | 1.03 |
| TSA (+) | 2.20 | 0.50 | 5.38 | 0.46 | --- | --- | 3.66 | 3.56 ± 0.15 | 1.10 |

a: fluorescence lifetime(s); b: amplitude(s); c: mean lifetime(s); d: average mean lifetime(s); e: reduced $\chi^2$ for the fit

### **Synthesis of Transition State Analogue (TSA):**
### (8-hydroxy-2-oxa-bicyclo[3.3.1]non-6-ene-3,5- dicarboxylic acid)

**Methyl 1-(2-methoxy-2-oxoethyl)cyclohex-3-ene-1-carboxylate (3):**

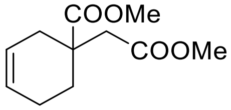


Dimethyl itaconate **1**(83.76 g, 529 mmol) and 3-sulfolene **2** (25.0 g, 211 mmol) in toluene (500 mL) were taken in 1000 mL PAR- reactor and heated to 90 °C and stirred for 24 h period (at which time no starting ester remained, TLC analysis). After cooling to rt, the solvent was evaporated, and the crude product was purified through silica gel column chromatography to yield corresponding Diels-Alder adduct **3** (35.92 g, 80%). $R_f$ = 0.5 (30% EtOAc-Hexane). $[\alpha]^{25}_{D}$= -0.29 (*c* 1.6, $CHCl_3$). IR υ max: 3027, 2965, 1740, 1511, 1445, 1016 $cm^{-1}$. $^1H$ NMR (400 MHz, $CDCl_3$): δ 5.70 – 5.60 (m, 2H), 3.70 (s, 3H), 3.65 (s, 3H), 2.10 – 1.76 (m, 8H) ppm. $^{13}C$ NMR (100 MHz, $CDCl_3$): δ 176.3, 171.6, 125.6, 124.1, 51.7, 51.4, 42.6, 40.1, 32.3, 29.2, 22.0 ppm. HRMS(ESI) *m/z* calculated for $C_{11}H_{16}O_4Na$ 235.0946, found 235.0939.

**2-(1-(Methoxycarbonyl)cyclohex-3-en-1-yl)acetic acid (4):**

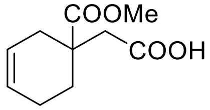


A solution of diester **3** (35.0 g, 165.0 mmol) in MeOH (293 mL) and NaOH (9.9 g, 247 mmol) and water (40 mL) was stirred at 21 °C for 6 days. The MeOH was evaporated, and the residue was diluted with water (100 mL), and the aqueous phase was washed with $CH_2Cl_2$. After acidification with concentrated HC1 to a pH of 2, product was extracted with $CH_2C1_2$ (3 x 200 mL), and finally evaporation of DCM afforded the monoacid **4** in 99% (32.36 g) yield as white needles: mp 62-65 °C. $[\alpha]^{25}_{D}$= +1.55 (*c* 0.28, $CHCl_3$). IR υ max: 3246, 3029, 2943, 1733, 1444, 1299, 1214 $cm^{-1}$. $^1H$ NMR (400 MHz, $CDCl_3$): δ 5.74 – 5.57 (m, 2H), 3.70 (s, 3H), 2.78 – 2.55 (m, 3H), 2.16 – 1.76 (m, 5H) ppm. $^{13}C$ NMR (100 MHz, $CDCl_3$): δ 177.2, 176.6, 125.8, 124.2, 52.2, 42.6, 39.9, 32.5, 29.3, 22.1 ppm. HRMS(ESI) *m/z* calculated for $C_{10}H_{14}O_4Na$ 221.0790, found 221.0785 $[M+H]^+$.

**Methyl 8-iodo-3-oxo-2-oxabicyclo[3.3.1]nonane-5-carboxylate (5).**

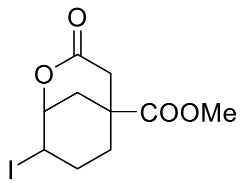

A solution of mono acid **4** (5.0 g, 25.2 mmol) and $NaHCO_3$ (4.23 g, 50.4 mmol), in water (400 mL) was treated with a solution of 6.4 g (25.2 mmol) of iodine and 25.1 g (151.1 mmol) of KI in water (900 mL). After being stirred for 11 h in the dark, the mixture was extracted several times with diethyl ether. The combined ether layers were washed with brine (200 mL), dried over anhydrous $MgSO_4$, and evaporated to give iodo lactone **5** (6.1 g, 75% yield) as a white solid. mp 92 °C. $R_f$ = 0.4 (30% EtOAc-Hexane). $[\alpha]^{25}_D$= +3.27 (*c* 0.3, $CHCl_3$). IR υ max: 2953, 2859, 1734, 1447, 1368, 1249, 1201 $cm^{-1}$. $^1H$ NMR (400 MHz, $CDCl_3$): δ 4.85 – 4.81 (m, 1H), 4.57 – 4.53 (m, 1H), 3.74 (s, 3H), 3.01 – 2.85 (m, 2H), 2.68 – 2.62 (m, 1H), 2.29 – 2.20 (m, 1H), 2.12 – 1.91 (m, 3H), 1.81 – 1.61 (m, 1H) ppm. $^{13}C$ NMR (100 MHz, $CDCl_3$): δ 174.2, 168.9, 77.9, 52.5, 40.5, 37.8, 29.2, 27.5, 26.2, 25.2 ppm. HRMS(ESI): *m/z* calculated for $C_{10}H_{14}O_4I$ 324.9937, found 324.9929 $(M+H)^+$.

**Methyl 3-oxo-2-oxabicyclo[3.3.1]non-7-ene-5-carboxylate (6).**

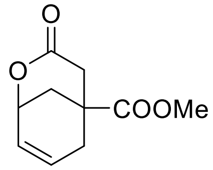


A solution of iodo lactone **5** (6.1 g, 18.8 mmol) and diazabicycloundecane (DBU) (4.22 mL, 28.1 mmol) in acetonitrile (100 mL) was heated at reflux for 5 h. The resulting mixture was partitioned between $CH_2C1_2$ and 1 N HC1, the aqueous layer was extracted with additional $CH_2C1_2$, and the combined organic layer was washed with brine, dried over anhydrous $MgSO_4$, evaporated, to give lactone **6** (3.22 g, 87% yield) as an yellow oil. $R_f$ = 0.4 (30% EtOAc-Hexane). $[\alpha]^{25}_D$= +1.21 (*c* 0.26, $CHCl_3$). IR υ max: 2944, 1724, 1441, 1364, 1249, 1193, 1064 $cm^{-1}$. $^1H$ NMR (400 MHz, $CDCl_3$): δ 6.09 – 5.91 (m, 2H), 4.88 – 4.81 (m, 1H), 3.74 (s, 3H), 3.14 – 3.01 (m, 1H), 2.66 – 2.51 (m, 2H), 2.39 – 2.37 (m, 2H), 2.13 – 1.75 (m, 1H) ppm. $^{13}C$ NMR (100 MHz, $CDCl_3$): δ 174.9, 170.1, 129.1, 126.0, 69.5, 62.4, 39.3, 38.3, 36.4, 30.5 ppm. HRMS(ESI): $[M+H]^+$ *m/z* calculated for $C_{10}H_{13}O_4$ 197.0814, found 197.0809.

**Methyl 3-hydroxy-2-oxabicyclo[3.3.1]non-7-ene-5-carboxylate (7).**

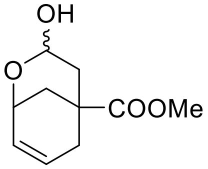


Lactone **6** (3.0 g, 15.0 mmol) was dissolved in THF (100 mL) under nitrogen atmosphere and cooled to -78 °C. Diisobutylaluminum hydride (DIBAL-H) (1.7 M solution in THF, 1.08 mL, 18.3 mmol) was added dropwise over 20 min and the reaction mixture was stirred at -78 °C for 1.5 h. The reaction was quenched by the addition of cold MeOH (10 mL) followed by 1 M HCl (25 mL)

and allowed to warm to room temperature. The mixture was then extracted with diethyl ether 50 mL and twice with $CH_2Cl_2$ (50 mL). The organic extracts were combined and washed with saturated $NaHCO_3$ solution until neutral. The organic layer was washed with 50 mL of water and 50 mL of brine and dried over $MgSO_4$, and the mixture was evaporated to give crude lactol. Immediate purification by column chromatography with 50% ethyl acetate/hexanes gave lactol **7** (2.18 g, 72% yield) as a white powder. Mp = 97 °C. $R_f$ = 0.3 (40% EtOAc-Hexane). IR υ max: 3436, 2955, 1731, 1443, 1249, 1184 $cm^{-1}$. $^1H$ NMR (400 MHz, $CDCl_3$): δ 6.07 – 5.99 (m, 1H), 5.86 – 5.77 (m, 1H), 5.27 – 5.16 (m, 1H), 4.46 – 4.38 (m, 1H), 3.70 (s, 3H), 3.07 – 2.98 (m, 1H), 2.61 – 2.48 (m, 1H), 2.32 – 2.23 (m, 1H), 2.17 – 2.00 (m, 2H), 1.87 – 1.75 (m, 1H), 1.71 – 1.57 (m, 1H) ppm. $^{13}C$ NMR (100 MHz, $CDCl_3$): δ 176.2, 131.6, 123.9, 89.5, 66.4, 52.1, 42.7, 40.6, 34.5, 32.1 ppm. MS(ESI): *m/z* 199 $[M+H]^+$.

**Methyl 3-methoxy-2-oxabicyclo[3.3.1]non-7-ene-5-carboxylate (8).**

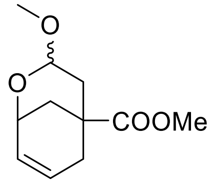


Lactol **7** (2.0 g, 10.0 mmol) was dissolved in methanol (100 mL) under $N_2$ and stirred in the presence of ten Nafion NR50 beads (ca. 30 mg) at 50 °C. After 15 h, the solution was filtered and the solvent was evaporated to give a clear oil. Chromatography of this compound with 40% ethyl acetate/hexane gave methyl acetal **8** (2.1 g, 97% yield) as a clear oil. $R_f$ = 0.6 (40% EtOAc-Hexane). IR υ max: 2952, 1732, 1443, 1394, 1248, 1180 $cm^{-1}$. $^1H$ NMR (400 MHz, $CDCl_3$): δ 6.08 – 6.01 (m, 1H), 5.87 – 5.79 (m, 1H), 4.84 (dd, $J$ = 3.2, 9.7 Hz, 1H), 4.49 – 4.42 (m, 1H), 3.71 (s, 3H), 3.45 (s, 3H), 2.59 – 2.49 (m, 1H), 2.32 – 2.22 (m, 1H), 2.18 – 2.10 (m, 1H), 1.97 – 1.89 (m, 1H), 1.86 – 1.77 (m, 1H), 1.76 – 1.68 (m, 1H) ppm. $^{13}C$ NMR (100 MHz, $CDCl_3$): δ 176.2, 131.4, 124.2, 96.4, 65.9, 56.1, 52.1, 40.3, 34.7, 32.2 ppm. HRMS(ESI) *m/z* calculated for $C_{11}H_{16}O_4Na$ 235.0946, found 235.0942.

**Methyl 3-cyano-2-oxabicyclo[3.3.1]non-7-ene-5-carboxylate (9).**

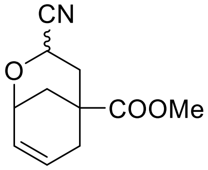


Methyl acetal **8** (2.0 g, 9.4 mmol) was dissolved in acetonitrile (80 mL) and cooled to 0 °C under $N_2$ with stirring. Zinc iodide (6.01 g, 18.8 mmol) was added quickly. After 5 min, trimethylsilyl cyanide (7.4 g, 75.2 mmol) was added drop wise over 10 min. and the solution was stirred for 2.5

h. Cold, saturated aq. $NaHCO_3$ (30 mL) was added and the mixture was warmed to room temperature. The mixture was extracted three times with $CH_2Cl_2$ (40 mL), and the organic layers were combined, washed with brine, and dried over $MgSO_4$. The solvent was evaporated to give an yellow oil, which was purified by chromatography using 33% ether/hexanes to give cyanohydrin ether **9** (1.6 g, 85% yield) as a clear oil, identified as an inseparable 1:1 exo/endo mixture. $R_f$ = 0.6 (40% EtOAc-Hexane). IR υ max: 2956, 1734, 1443, 1249, 1191, 1124, 1083 $cm^{-1}$. $^1H$ NMR (400 MHz, $CDCl_3$): δ 6.28 – 6.19 (m, 2H), 6.03 – 5.96 (m, 1H), 5.77 – 5.72 (m, 1H), 4.92 – 4.84 (m, 1H), 4.77 – 4.69 (m, 1H), 4.54 – 4.40 (m, 2H), 3.73 (s, 3H), 3.72 (s, 3H), 2.69 – 2.55 (m, 3H), 2.39 – 2.25 (m, 2H), 2.21 – 2.00 (m, 5H), 1.93 – 1.84 (m, 2H), 1.64 – 1.59 (m, 1H) ppm. $^{13}C$ NMR (100 MHz, $CDCl_3$): δ 175.8, 175.2, 135.2, 133.8, 124.0, 122.2, 119.8, 118.3, 66.7, 65.8, 57.1, 56.9, 52.4, 38.7, 38.2, 37.5, 35.2, 34.1, 31.6, 31.4 ppm. $C_{11}H_{13}NO_3$, 207 MS(ESI): *m/z* 208 $[M+H]^+$.

**Methyl (1*R**, 2*S**, 4*R**, 6*S**, 8*R***S**)-8-cyano-3,9-dioxatricyclo[4.3.1.0$^{2,4}$]decane-6-carboxylate (10).**

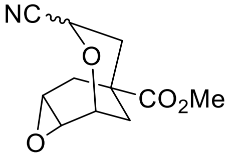


A solution of cyano olefin **9** (1.5 g, 7.0 mmol) and *m*-chloroperbenzoic acid (3.04 g, 17 mmol) in $CH_2C1_2$ (50 mL) was heated at reflux for 24 h. A solution of $K_2SO_3$ (1.0 g) in $H_2O$ (12 mL) was added, and the mixture was stirred overnight. The resulting suspension was washed with saturated $NaHCO_3$, 2 N NaOH, $H_2O$, and finally brine, and the organic layer was dried over $MgSO_4$, and evaporated to give crude epoxide **10** as a white solid. Chromatographic purification of this material (2:l hexanes/$Et_2O$) afforded epoxide (1.56 g, 97% yield) as a 1:l mixture. $R_f$ = 0.7 (40% EtOAc-Hexane). IR υ max: 3008, 2953, 1734, 1443, 1367, 1249, 1205 $cm^{-1}$. $^1H$ NMR (400 MHz, $CDCl_3$): δ 4.92 – 4.88 (m, 1H), 4.56 – 4.52 (m, 1H), 3.71 (s, 3H), 3.52 – 3.48 (m, 1H), 3.39 – 3.36 (m, 1H), 2.57 – 2.50 (m, 1H), 2.39 – 2.28 (m, 2H), 2.12 – 2.01 (m, 2H), 1.70 – 1.55 (m, 1H) ppm. $^{13}C$ NMR (100 MHz, $CDCl_3$): δ 175.0, 119.7, 68.7, 57.7, 53.2, 52.5, 50.6, 36.1, 35.6, 30.2, 26.8 ppm. $C_{11}H_{13}NO_4$, MS(ESI): *m/z* 246 $[M+Na]^+$.

**Methyl (8-exo)-3-cyano-8-hydroxy-2-oxabicyclo[3.3.l]non6-ene-5-carboxylate (11).**

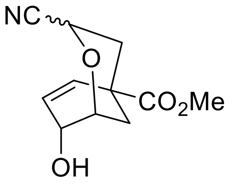


To the stirring solution of the epoxide **10** (1.0 g, 4.4 mmol) and triphenylphosphine (691 mg, 2.6 mmol) in acetonitrile (10 mL) was added trimethylsilyl bromide (TMSBr) (1.3 g, 8.8 mmol) dropwise at 0 °C. After 0.5 h, the mixture was warmed to 20 °C and stirred for an additional 1.5 h.

1,5-diazabicyclo[5.4.0]undec-5-ene (DBU) (1.67 g, 11.0 mmol) was then added dropwise and the mixture was heated at reflux for 17 h. After evaporation of the mixture, the residue was triturated with $Et_2O$, leaving a brown precipitate of DBU.HBr. The supernatant was filtered through Celite and evaporated, and the residue was dissolved in THF (4.5 mL). Concentrated HCl was added dropwise until a pH of 1 was attained. The mixture was evaporated, the residue was dissolved in $CH_2Cl_2$, and the solution was washed with saturated $NaHCO_3$, and brine, dried ($Na_2SO_4$), and evaporated. Chromatography of the resulting yellow oil (4:1 ether/hexanes) gave 770 mg (77% yield) of allylic alcohol **11** as a 1:l mixture. $R_f$ = 0.4 (50% EtOAc-Hexane). IR υ max: 3438, 2957, 1729, 1441, 1399, 1252, 1185 $cm^{-1}$. $^1H$ NMR (400 MHz, $CDCl_3$): δ 6.27 – 6.18 (m, 3H), 6.12 – 6.08 (m, 1H), 4.89 (d, 1H, $J$ = 7.4 Hz), 4.52 (dd, $J$ = 3.2, 12.2 Hz, 1H), 4.43 – 4.41 (m, 1H), 4.24 – 4.18 (m, 2H), 4.09-4.07 (m, 1H), 3.77 (s, 3H), 3.78 (s, 3H), 2.34 – 2.29 (m, 1H), 2.25 – 2.14 (m, 2H), 2.05 – 2.03 (m, 2H), 1.97-1.89 (m, 3H), 1.82-1.75 (m, 2H), ppm. $^{13}C$ NMR (100 MHz, $CDCl_3$): δ $^{13}C$ NMR (126 MHz, $CDCl_3$) δ 174.13, 173.70, 132.67, 131.65, 130.36, 119.07, 117.80, 73.30, 73.18, 65.31, 59.14, 58.64, 52.72, 52.69, 41.04, 39.41, 34.28, 32.35, 28.37, 28.16. ppm. MS(ESI): calculated for $C_{11}H_{13}NO_4$: *m/z* 223 $[M]^+$

**(3–exo,8-exo)-8-Hydroxy-2-oxabicyclo[3.3.1]non-6-ene-3,5-dicarboxylic acid and the (3–endo,8-exo) isomer (12 & 13).**

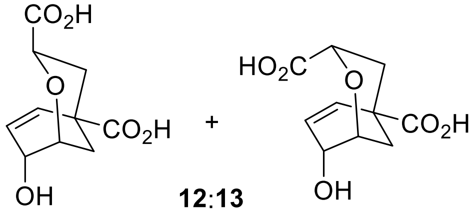


**12**:**13**
3:1

The solution of nitrile ester **11** (0.2 g, 0.89 mmol) and 85% KOH (0.120 g, 2.1 mmol) in water (6 mL) was heated at reflux for 2 h. The mixture was acidified to pH 2 with 2 N HCl and concentrated under reduced pressure to give the crude mixture of diacids. The di acid residue was triturated with 4 portions of acetone. The combined acetone solutions were concentrated under reduced pressure and dried under vacuum over $P_2O_5$ to give the hygroscopic diacids **12** & **13** (0.19 g, 97%) as 3:1 mixture of the exo:endo isomers (isomer ratio is based on NMR spectrum). Rf = 0.7 (40% EtOAc-Hexane). IR υ max: 3479, 3021, 2955, 1797, 1435, 1215 $cm^{-1}$. $^1H$ NMR (500 MHz, $CD_3OD$) for Exo isomer: δ 6.21-6.14 (m, 2H), 4.25 (dd, 1H, $J$ = 3.3, 12.2 Hz), 4.13 (brs, 1H), 4.01 (s, 1H), 2.05 – 1.81 (m, 4H) ppm. Endo isomer: $^1H$ NMR (500 MHz, MeOD) δ 6.05 (dd, J = 10.1, 0.9 Hz, 2H), 5.88 – 5.81 (m, 2H), 4.43 (dd, J = 7.6, 2.2 Hz, 2H), 4.33 (ddd, J = 7.0, 3.5, 2.5 Hz, 2H), 4.13 (brs, 1H). $^{13}C$ NMR (125 MHz, $CD_3OD$): Exo: δ 177.7, 175.3, 133.0, 132.4, 130.9, 130.8, 79.4, 74.8, 74.4, 70.3, 69.4, 56.9, 66.0, 42.6, 41.2, 35.2, 33.0, 29.9, 28.8 ppm. MS(ESI): *m/z* 246 $[M+NH_4]^+$.

A slightly modified strategy in comparison to known route[1] was followed to synthesize **12** and **13** as a mixture of diacids in 3:1 ratio of exo:endo isomers. Accordingly, the synthesis of oxabicyclo[3.3.1]nonene **12** (exo) and **13** (endo) as a 3:1 mixture is outlined below in the scheme. The Diels-Alder reaction of butadiene generated insitu from 3-sulfolene(butadiene cyclic sulfone) **1** with dimethyl itaconate **2** afforded the adduct **3** which on hydrolysis (less hindered ester hydrolysis) with NaOH furnished mono acid **4**. Iodolactonization of monoester acid **4** with KI and $I_2$ provided iodide **5**, which was further subjected to an elimination reaction with DBU to provide bicyclic olefin **6** as a 1:1 mixture of epimers**.** Treatment of lactone **6** with DIBAL-H provided lactol which was further protected as hemiacetal **8** with MeOH in presence of nafion beads. Cyanation of hemiacetal **8** was achieved with TMSCN in presence of $ZnI_2$ to provide the corresponding cyano compound **9** in decent yields. The olefin **9** upon epoxidation with mCPBA furnished **10**. Conversion of **10** to **11** was achieved in one pot reaction by treatment of **10** with bromotrimethylsilane, and triphenylphosphine (catalyst) to form the silyl bromohydrin followed by subsequent elimination reaction upon treatment with diazabicycloundecene (DBU). Hydrolysis of the mixture of epimeric nitrile ester **11** was achieved with KOH to furnish the diacids **12** (exo) and **13** (endo). The exo and endo ratio and configurations of **12** and **13** were identified based on $^1H$ NMR coupling constants. No further attempts were made to separate the mixture and the 3:1 mixture was used directly for the biological studies.

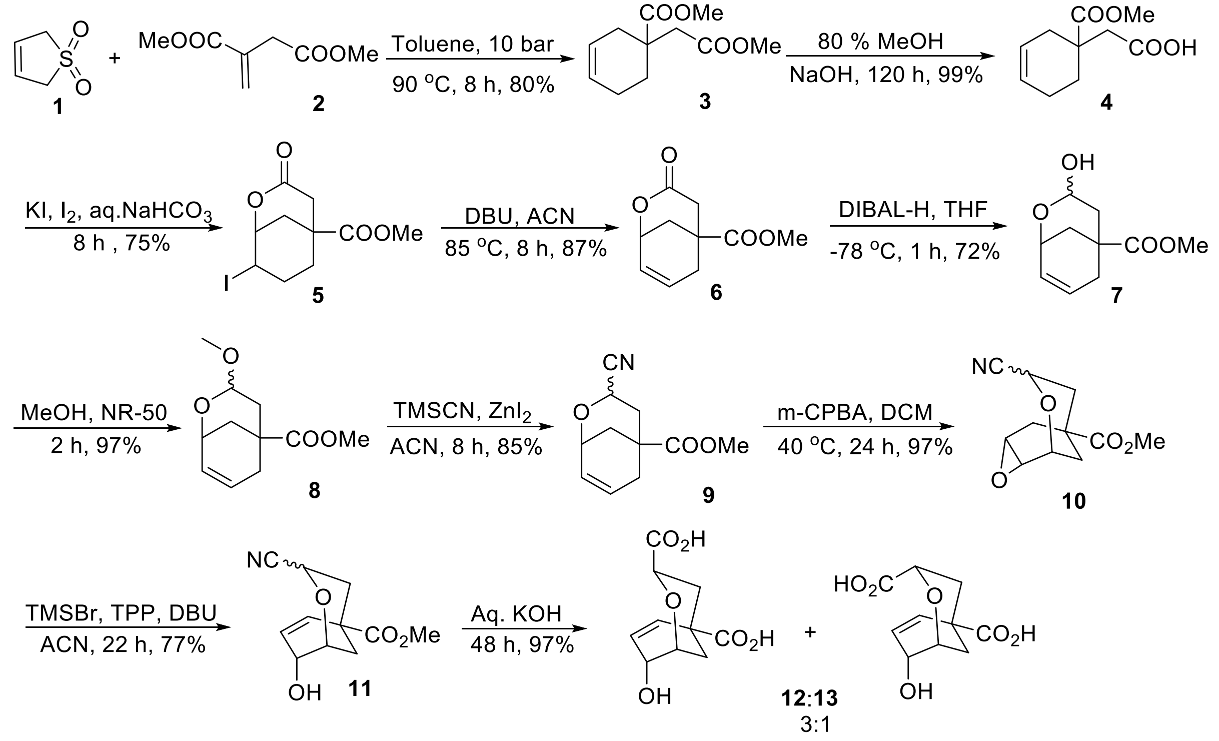



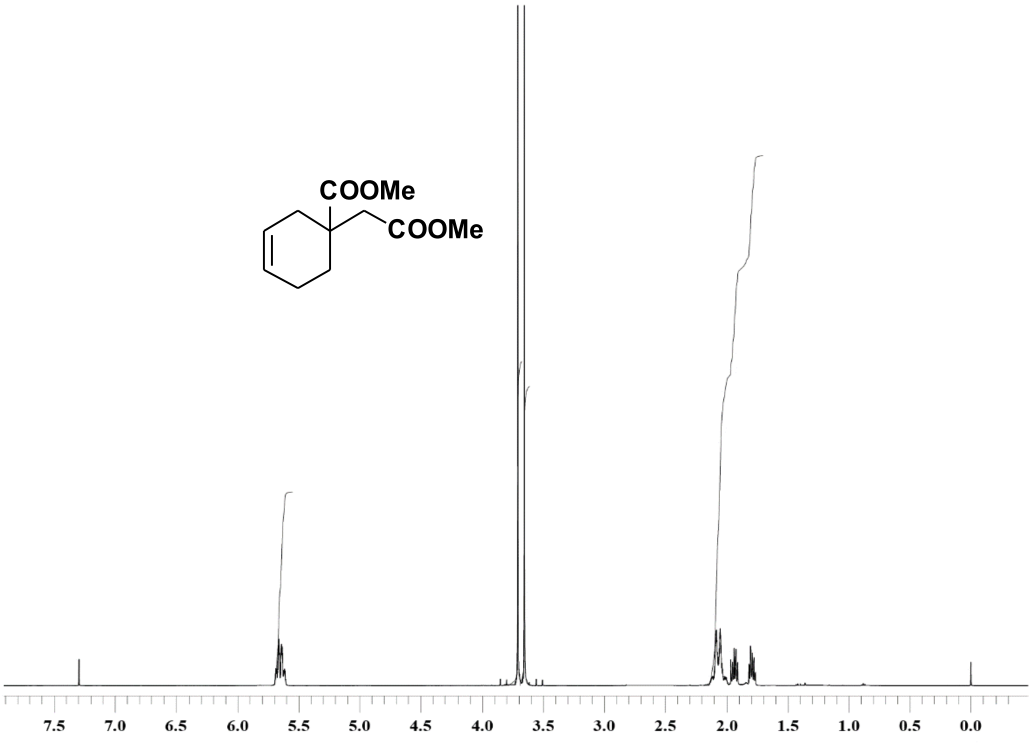


Compound **3** $^{1}$H NMR, 400 MHz, $CDCl_3$

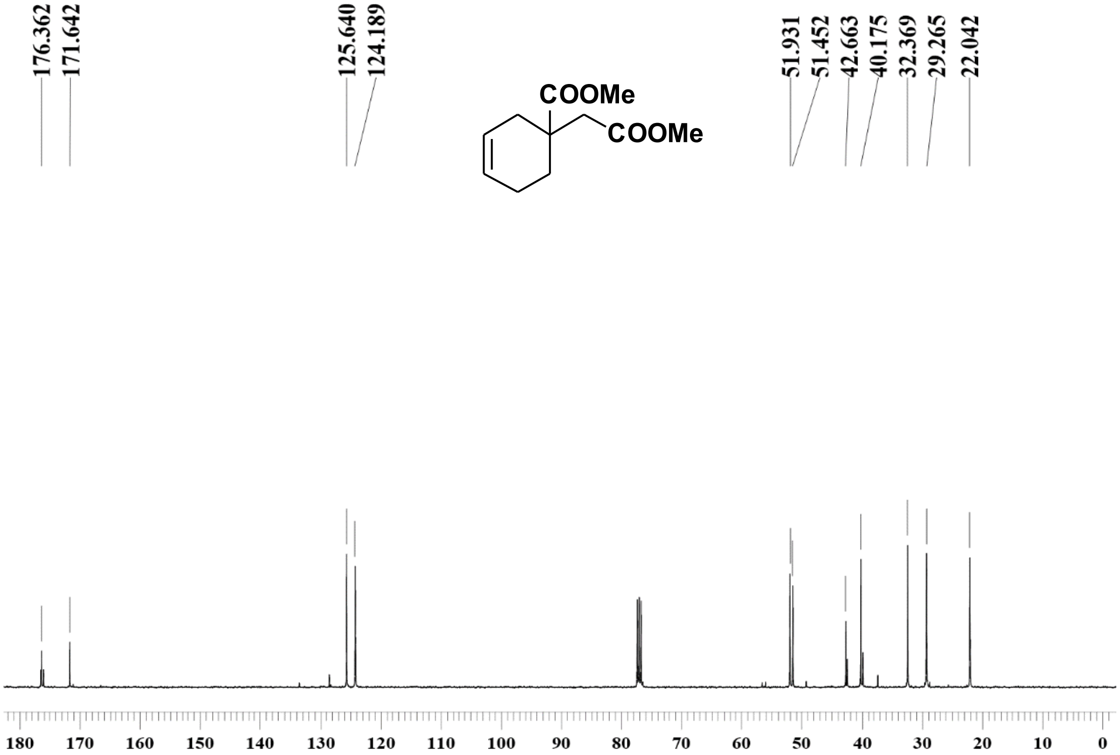


Compound **3** $^{13}$C NMR, 100 MHz, $CDCl_3$

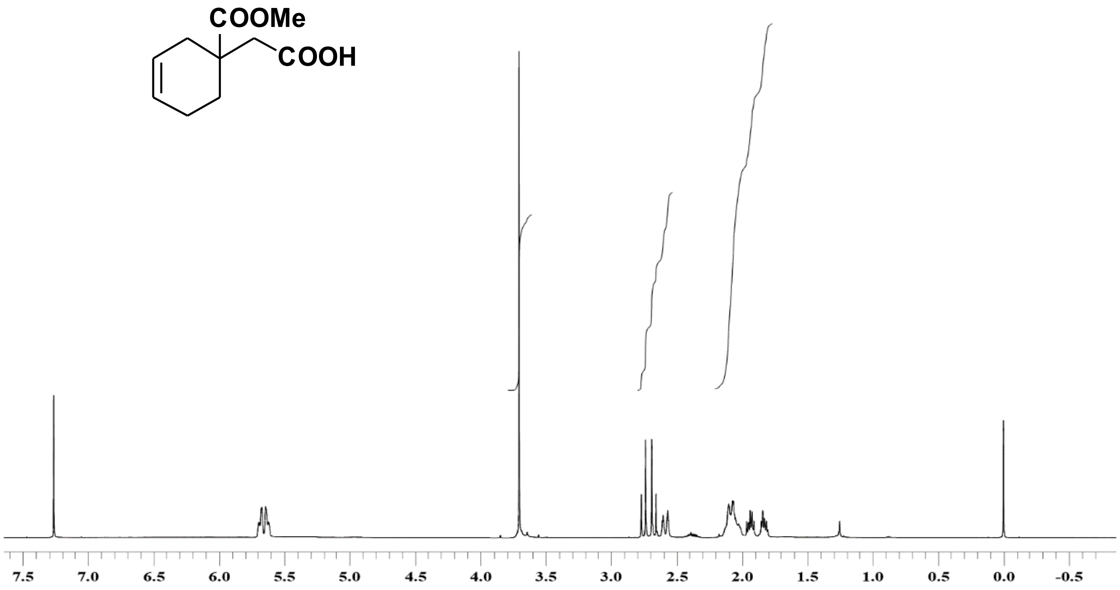


Compound **4** -$^{1}$H NMR, 400 MHz, $CDCl_3$

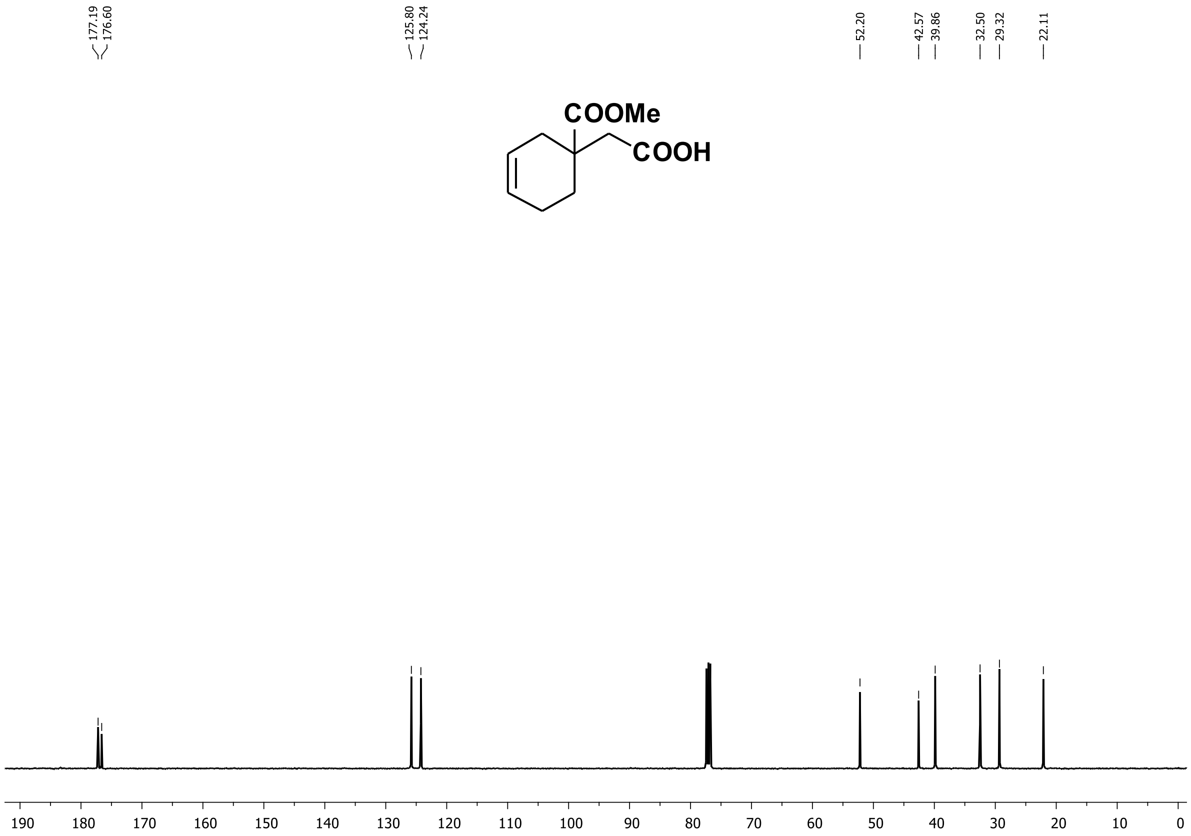


Compound **4** $^{13}$C NMR, 100 MHz, $CDCl_3$

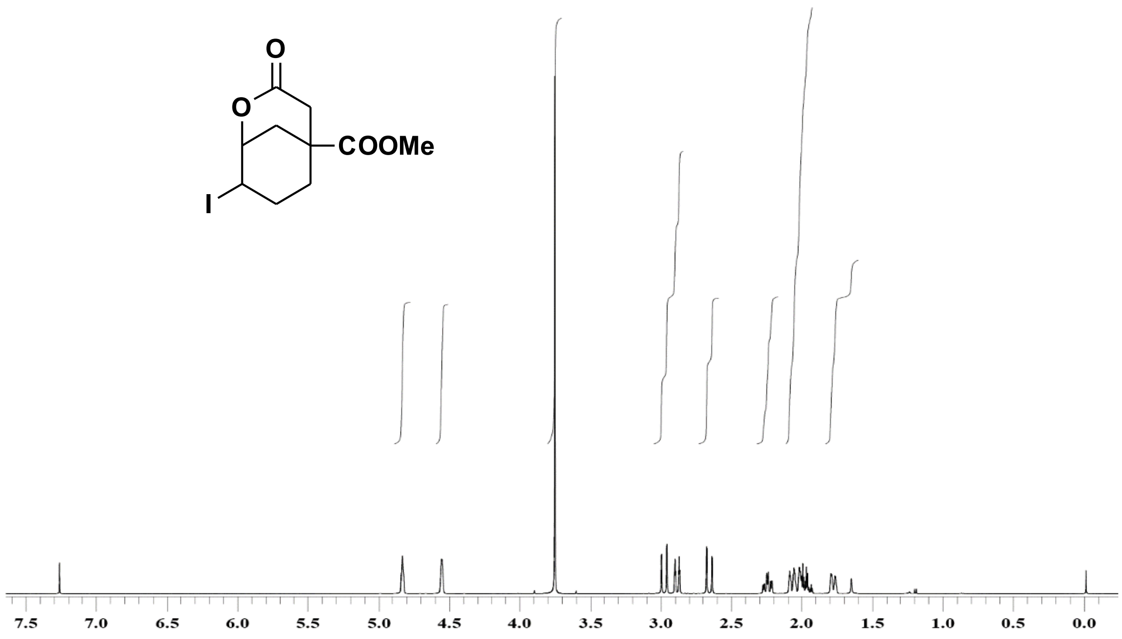


Compound **5** $^{1}$H NMR, 400 MHz, $CDCl_3$

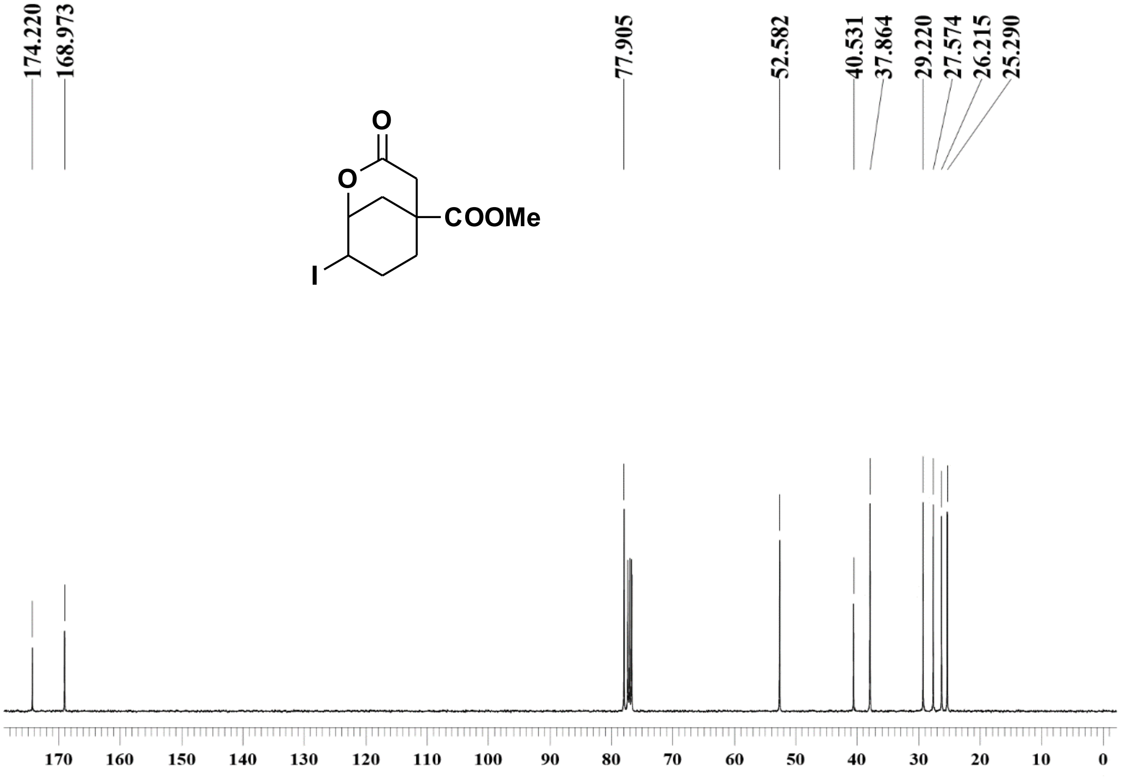


Compound **5** $^{13}$C NMR, 100 MHz, $CDCl_3$

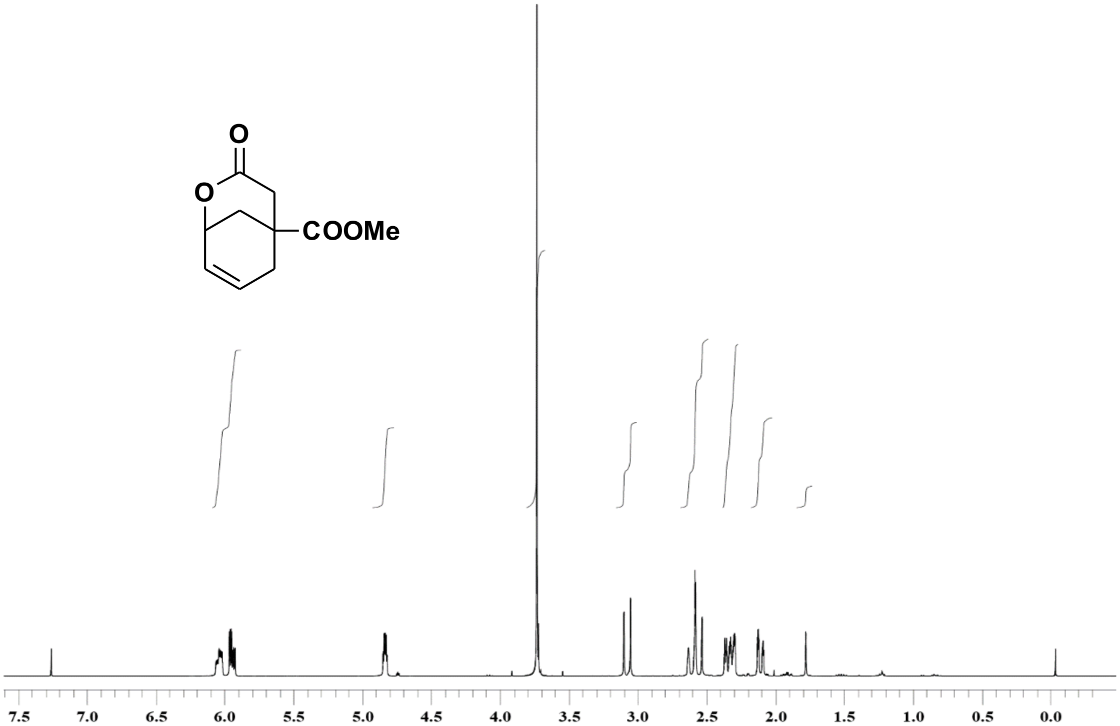


Compound **6** $^{1}$H NMR, 400 MHz, $CDCl_3$

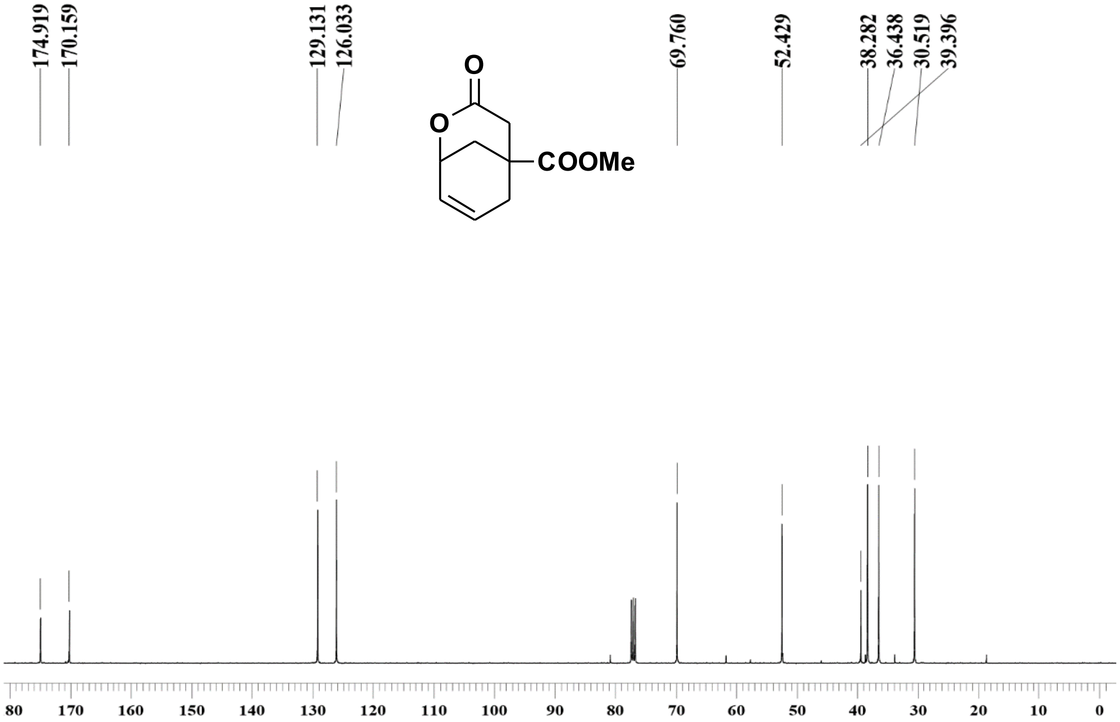


Compound **6** $^{13}$C NMR, 100 MHz, $CDCl_3$

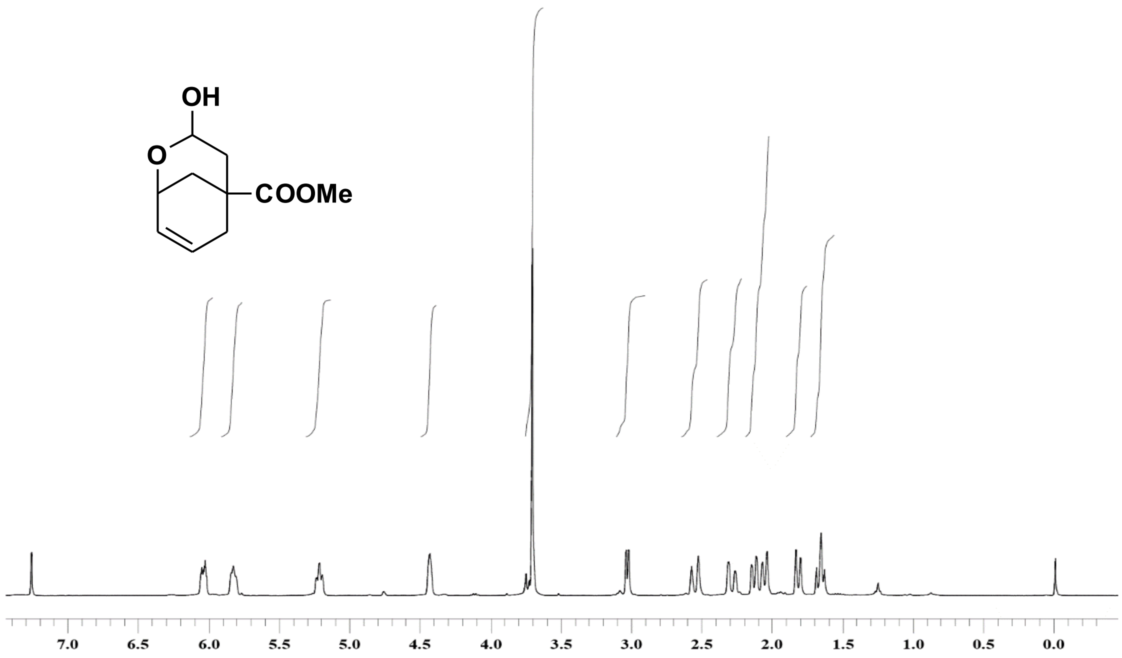


Compound **7** $^{1}$H NMR, 400 MHz, $CDCl_3$

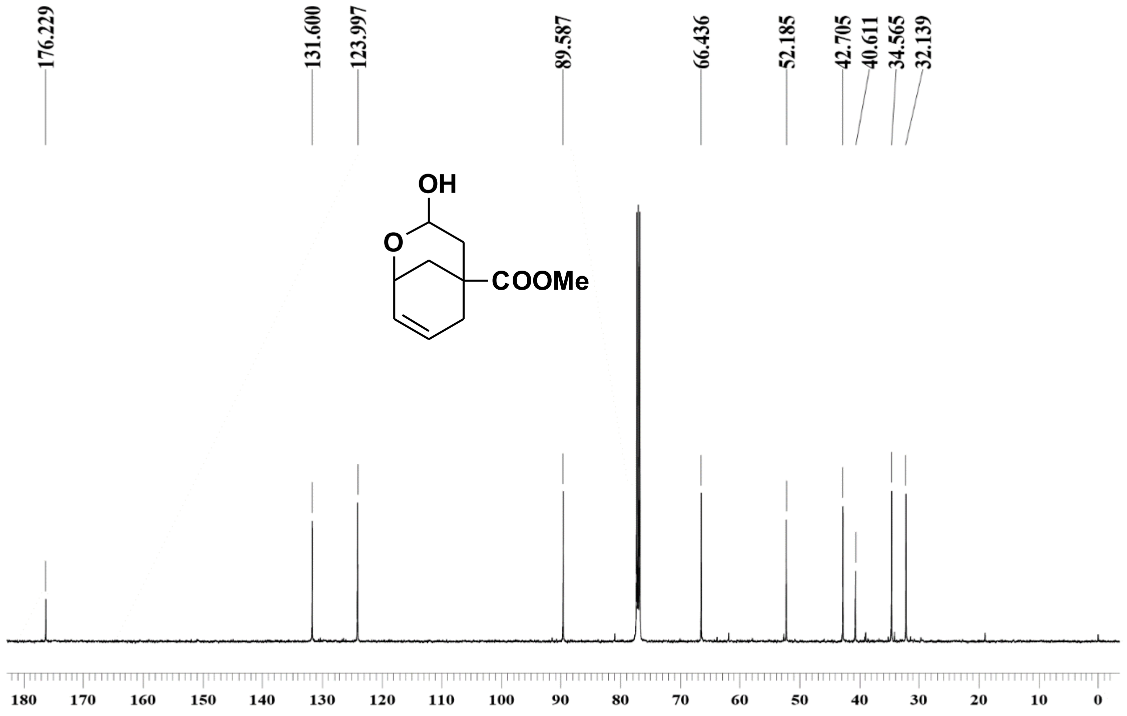


Compound **7** $^{13}C$ NMR, 100 MHz, $CDCl_3$

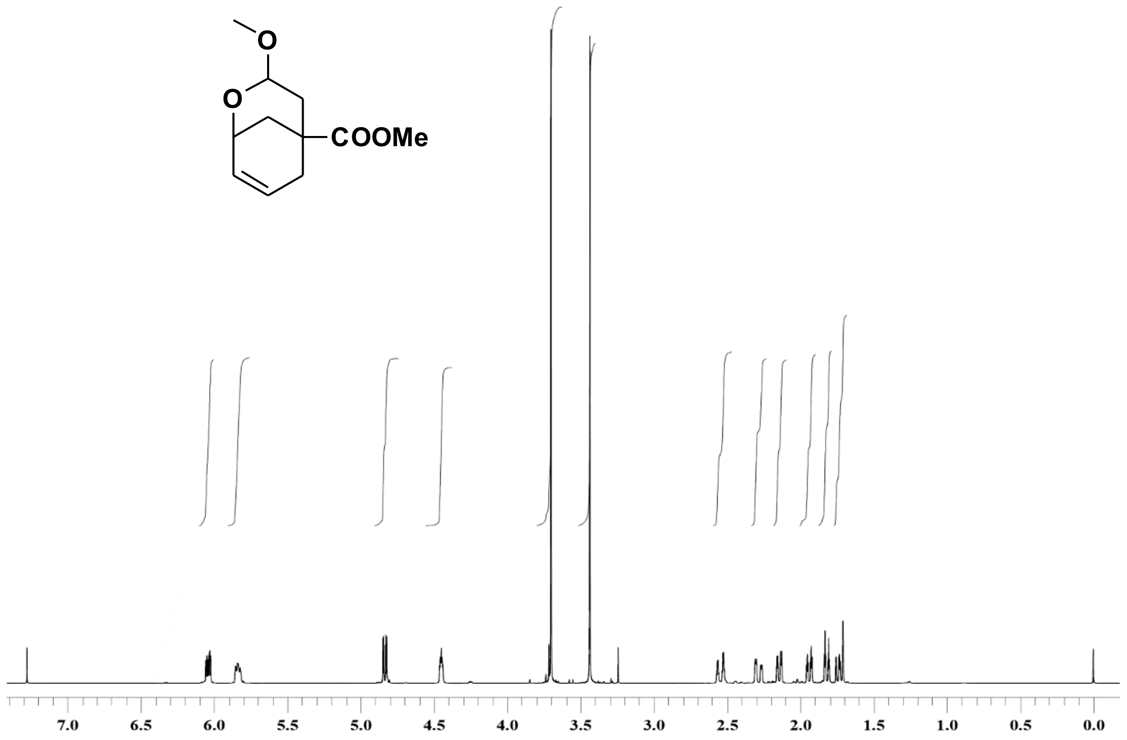

Compound **8** $^{1}$H NMR, 400 MHz, CDCl$_3$

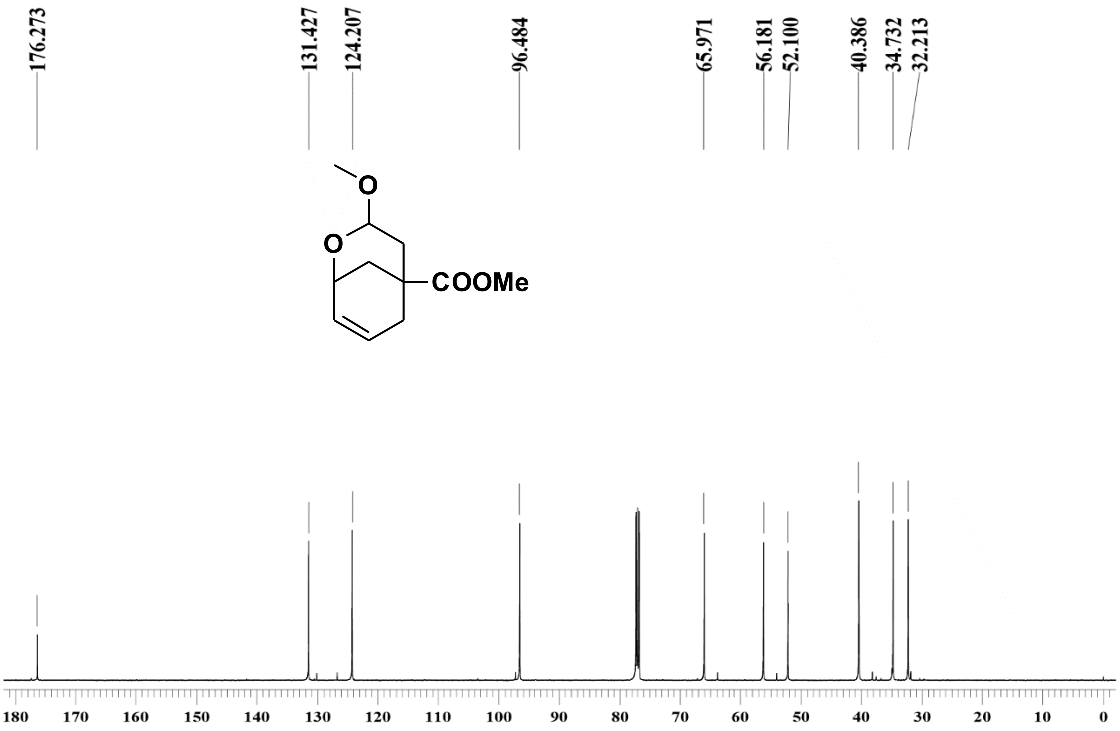


Compound **8** $^{13}$C NMR, 100 MHz, CDCl$_3$

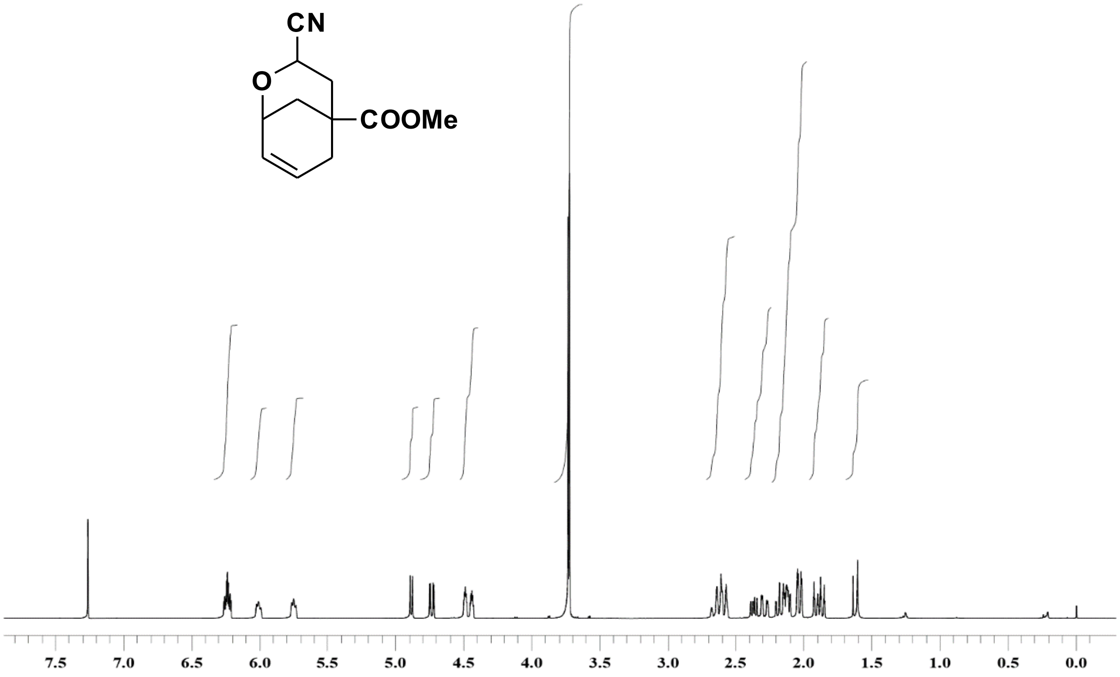


Compound **9** $^{1}H$ NMR, 400 MHz, $CDCl_3$

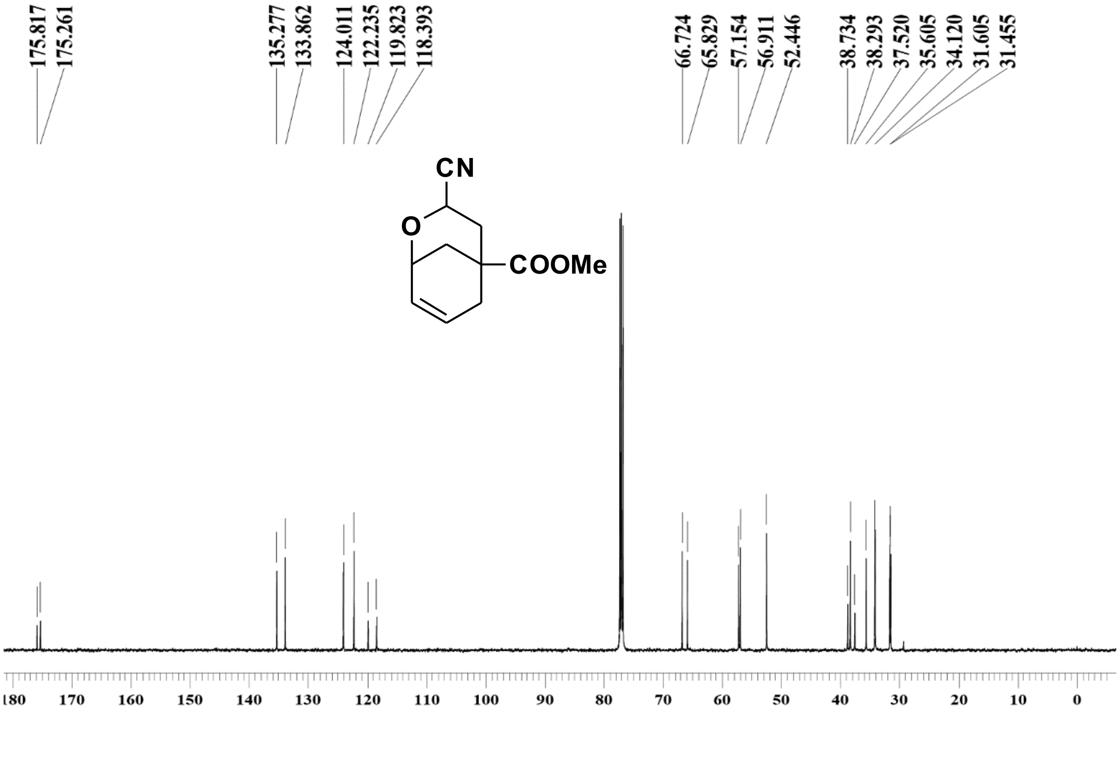


Compound **9** $^{13}C$ NMR, 100 MHz, $CDCl_3$

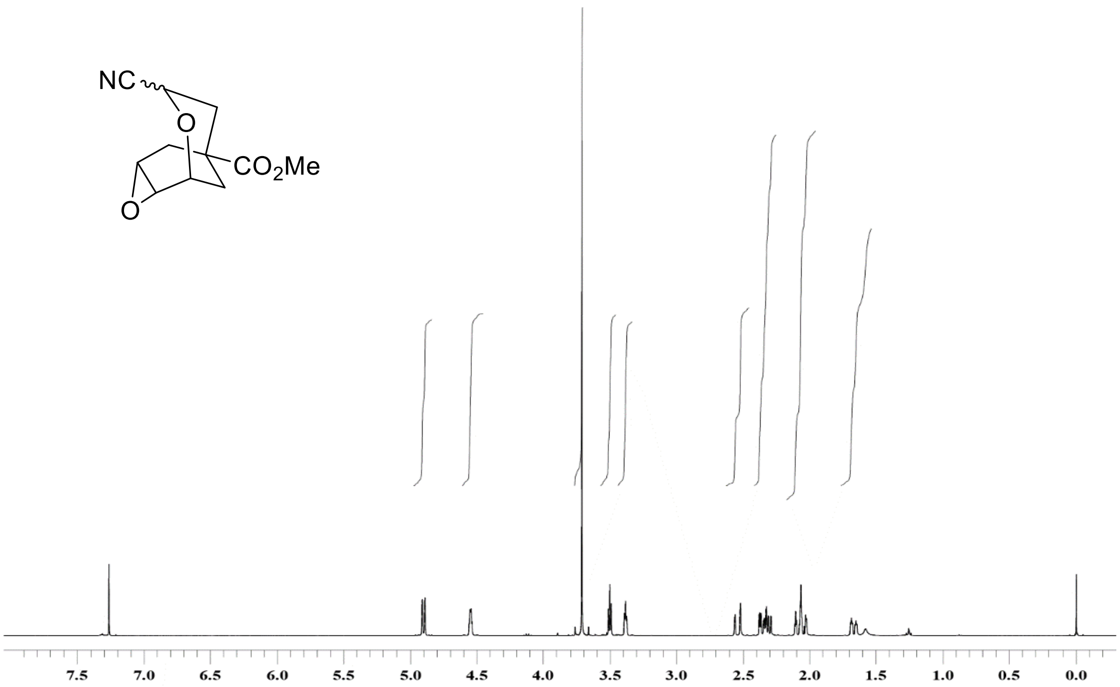


Compound **10** $^{1}$H NMR, 400 MHz, $CDCl_3$

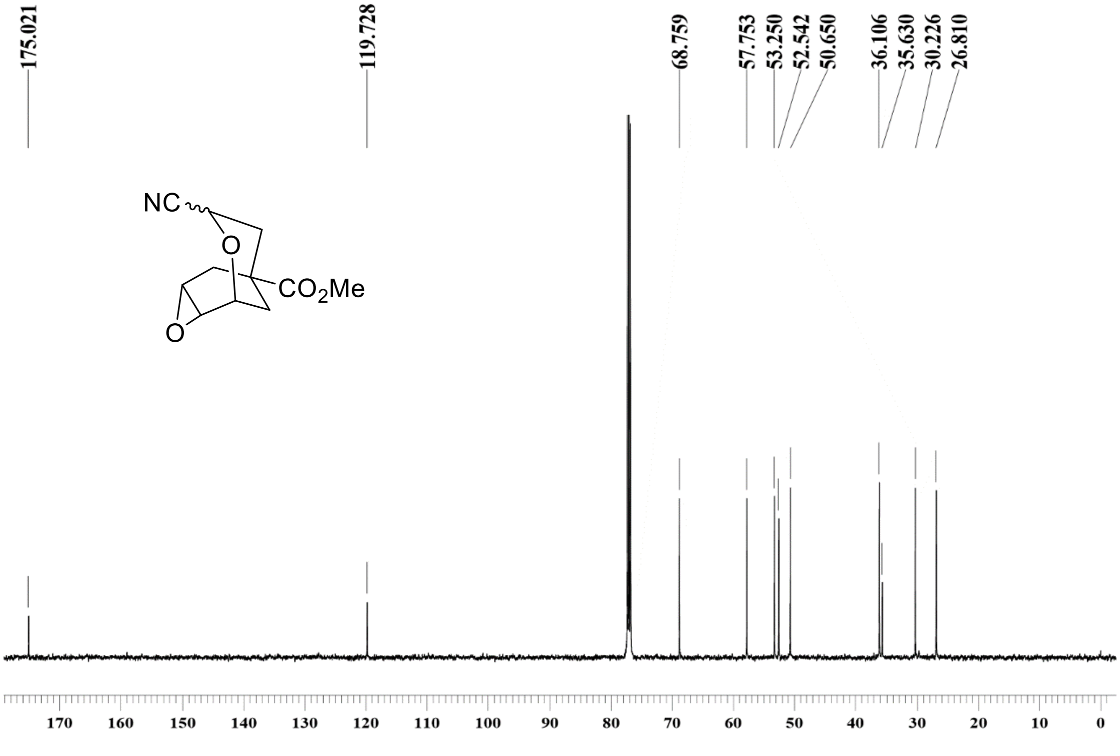


Compound **10** $^{13}$C NMR, 100 MHz, $CDCl_3$

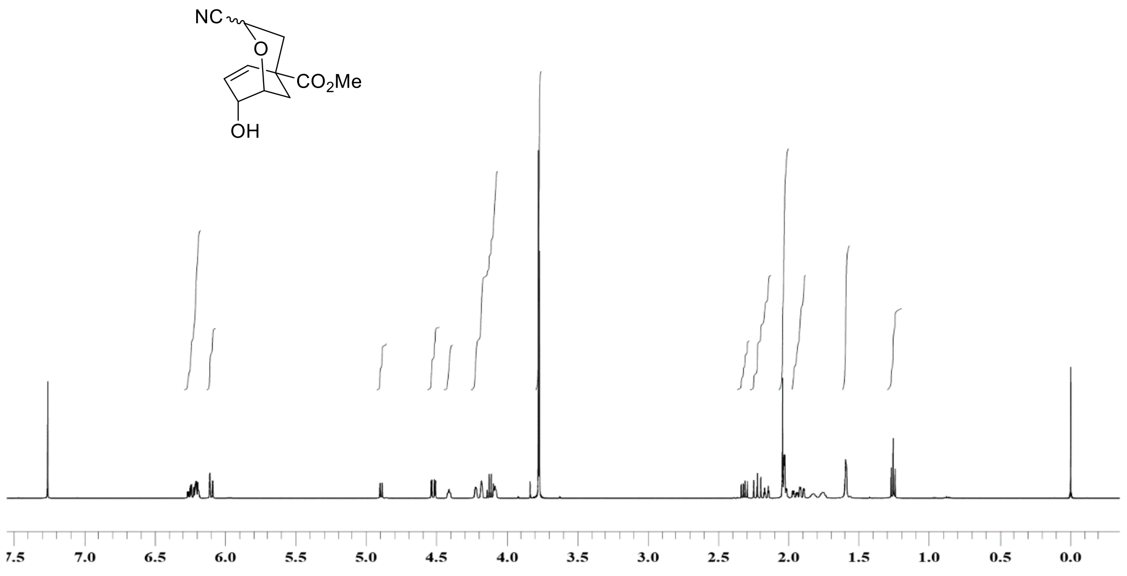

Compound **11** $^{1}H$ NMR, 400 MHz, $CDCl_3$

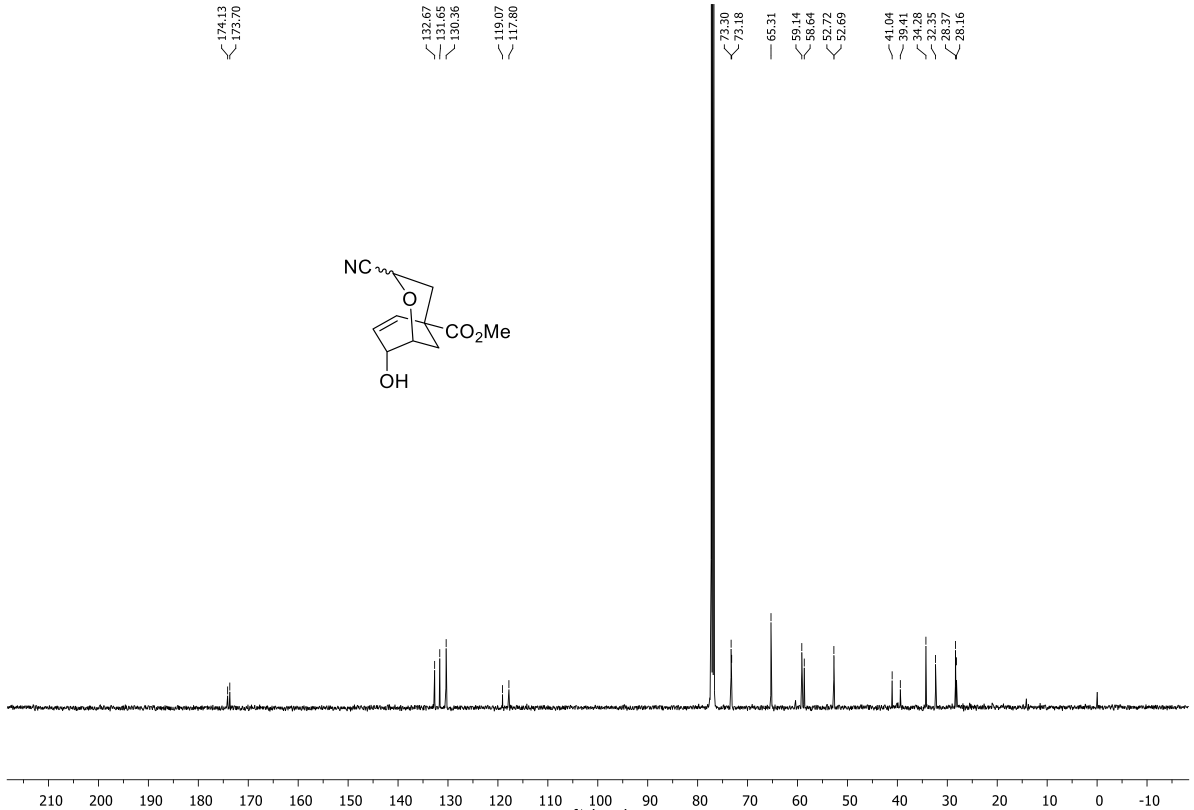

Compound **11** $^{13}C$ NMR, 100 MHz, $CDCl_3$

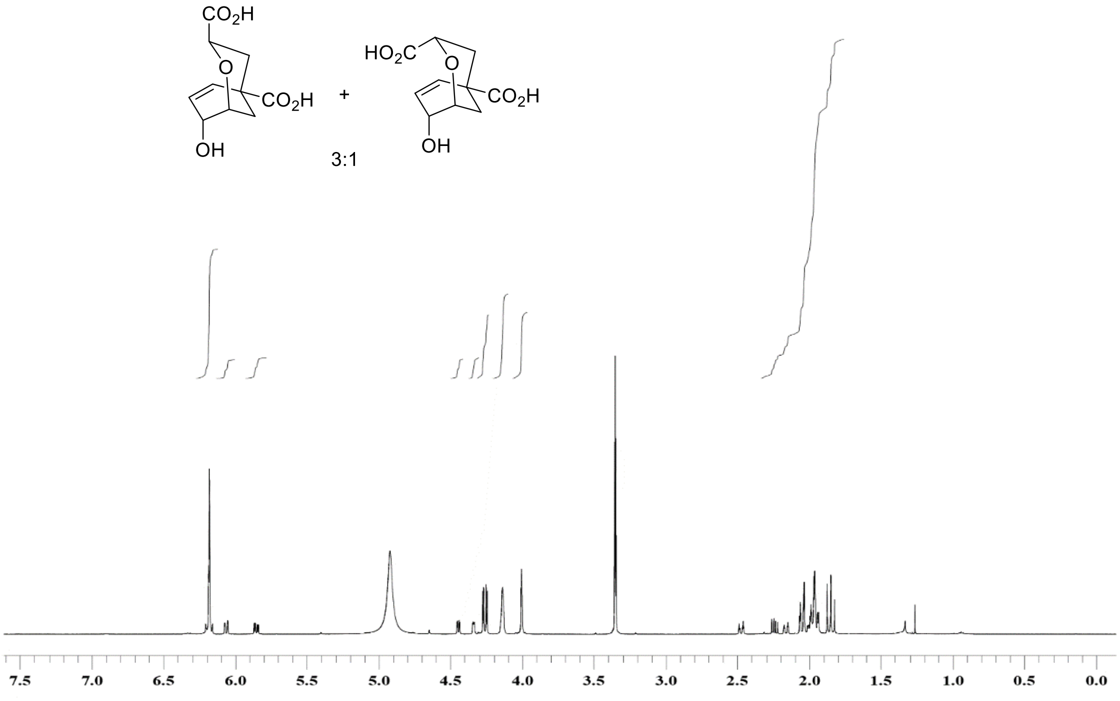


[1]H NMR for mixture of **12** & **13**, 500 MHz, $CD_3OD$

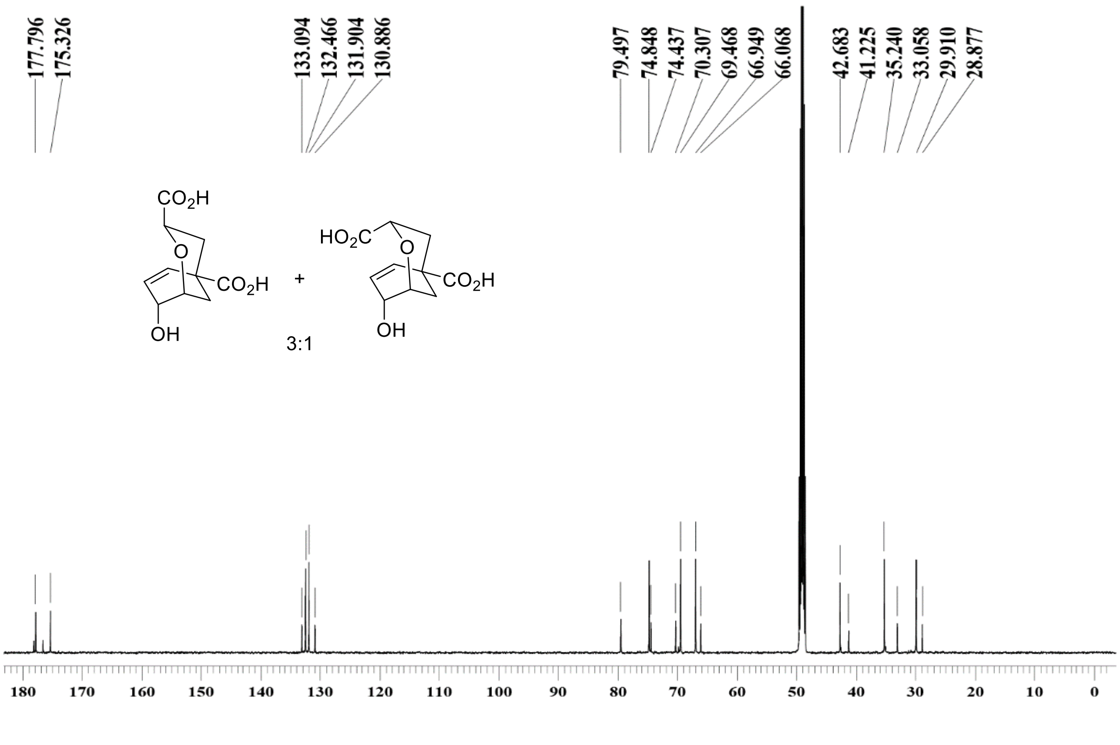


[13]C NMR for mixture of **12** & **13**, 125 MHz, $CD_3OD$
